# iSanXoT: a Standalone Application for the Integrative Analysis of Mass Spectrometry-Based Quantitative Proteomics Data

**DOI:** 10.1101/2023.08.25.554773

**Authors:** Jose Manuel Rodríguez, Inmaculada Jorge, Rafael Barrero-Rodríguez, Ricardo Magni, Estefanía Núñez, Andrea Laguillo, Cristina A. Devesa, Juan A. López, Emilio Camafeita, Jesús Vázquez

**Affiliations:** Proteomics Unit, Centro Nacional de Investigaciones Cardiovasculares Carlos III (CNIC), Madrid 28029, Spain; Laboratory of Cardiovascular Proteomics. Centro Nacional de Investigaciones Cardiovasculares (CNIC), Madrid 28029, Spain; CIBER de Enfermedades Cardiovasculares (CIBERCV), Madrid, Spain

**Keywords:** Mass spectrometry, Quantitative proteomics, Proteomics pipeline, Generic integration algorithm, WSPP model, protein coordination

## Abstract

Numerous bioinformatics tools currently exist to perform quantitative analysis of proteomics experiments. The majority of these tools apply diverse statistical models to assign a quantitative protein value from the mass-spectrometry information. Here we present iSanXoT, a standalone application that allows integrative analysis of quantitative proteomics data. iSanXoT processes relative abundances between MS signals and integrates them sequentially to upper levels using our previously published Generic Integration Algorithm (GIA). iSanXoT offers unique capabilities that complement conventional quantitative softwares, including statistical weighting and independent modeling of error distributions in each integration, aggregation of technical or biological replicates, quantification of posttranslational modifications or analysis of coordinated protein behavior. iSanXoT is a standalone, user-friendly application which accepts output from widespread proteomics pipelines and enables free construction of quantification workflows and fully customizable reports than can be reused across different projects or shared among users. Diverse integrative workflows constructed using GIA for the analysis of high-throughput quantitative proteomics experiments have been successfully applied in numerous publications. iSanXoT has been tested with the main operating systems. Download links for the corresponding distributions are available at https://github.com/CNIC-Proteomics/iSanXoT/releases.

## Introduction

Numerous bioinformatic packages, such as MaxQuant [1], FragPipe [2] or Trans Proteomics Pipeline [3], currently exist to perform quantitative analysis of proteins from mass spectrometry (MS)-based proteomics data. The majority of them process the raw data (i.e. MS signal) to assign a quantitative value for each protein in each sample. The package SanXoT [4] was designed to process relative abundances between MS signals, which were sequentially integrated to obtain weighted averages of protein ratios by the iterative application of the Generic Integration Algorithm (GIA) [5]. This approach exploits the stability of intensity ratios within the same scan (spectrum) in stable isotope labeling experiments and allows separate and accurate modeling of the different error sources at the scan, peptide and protein levels, as shown in the original WSPP model [6]. The SanXoT suite also allows to perform some types of quantitative analysis that are straightforward applications of the GIA algorithm, such as the integration of protein values across experiments performed using different techniques [6], the analysis of coordinated protein behavior using the Systems Biology Triangle (SBT model) [5] and the quantitative analysis of post-translational modifications [7, 8]. SanXoT has been used in more than 60 published quantitative proteomics studies (see for instance [9–18]).

The SanXoT package was originally developed for bioinformaticians and needed the modular assembly of a series of scripts to construct quantitative workflows, which once mounted could not be easily reused across different experiments. Besides, tools were lacking for the preparation of result tables and for the adaptation of data from other software packages. Hence, in spite of its potential, these complications, pointed out by SanXoT users, have hindered the widespread use of this package by the proteomics community. Here we present iSanXoT, an open-source application that incorporates the set of SanXoT programs within a user-friendly interface without losing its flexibility to construct limitless, fully customizable user-designed quantitative workflows. Once mounted, the workflows can be stored and easily reused for the analysis of new datasets. iSanXoT includes a reporting module that allows to extract the quantitative information and generate user-customizable result tables gathering the desired quantitative data from the specified analysis levels. Finally, iSanXoT also includes a general adaptor to process quantitative data obtained from other widely-used software tools, such as MaxQuant, Fragpipe or Proteome Discoverer (Figure 1).

**Figure 1.**
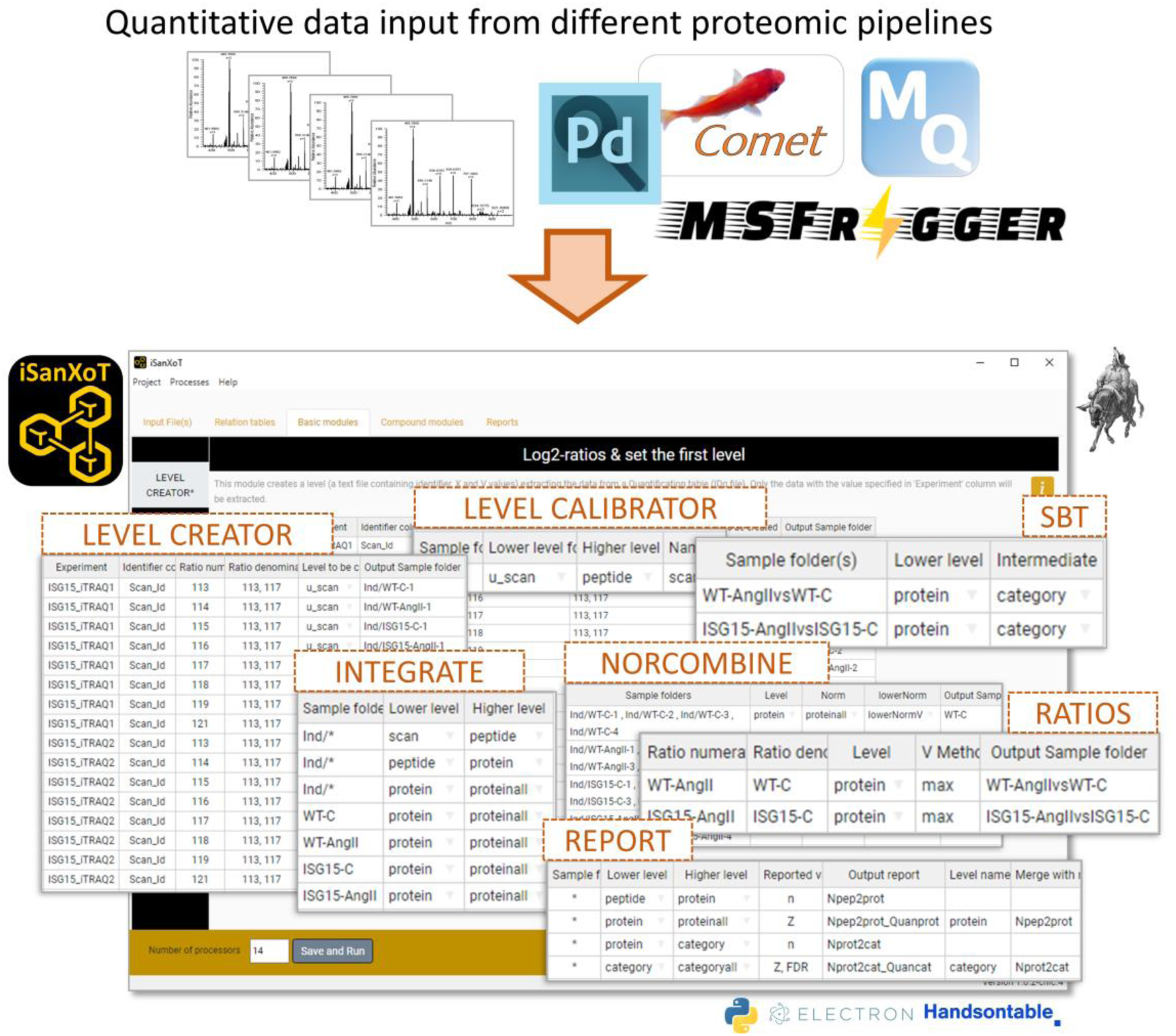
Scheme of the modular design of iSanXoT. iSanXoT accepts quantitative data from widely-used proteomics pipelines. The set of modules needed to run the quantitative workflows is invoked within a user-friendly interface based on task tables, allowing fully customizable user-designed quantitative workflows.

## Materials and methods

iSanXoT uses the GIA algorithm [5] to sequentially integrate the quantitative information (relative abundance) at increasing levels allowing the construction of fully customizable workflows that can be adapted to a wide range of needs. iSanXoT houses several modules built upon the programs included in the SanXoT package. In general, the workflows start by adapting the output from other sofware packages to be used with iSanXoT. This is followed by the sequential integration of the quantitative information at increasing levels and the generation of reports containing the desired information, as specified by the user through an user-friendly, task table interface. The interface contains several basic modules which performs separate steps, and also compound modules that contain commonly used workflows ready to be used. In addition, iSanXoT allows newly created workflows to be easily reused by the same or other users. In this work we include four representative workflows that not only illustrate the flexibility of the package, but that may also be reused or adapted by any user. The Supplementary Information includes very detailed information about how to use these workflows and how to use quantitative data from some popular proteomics pipelines.

### Input Data Adapter

iSanXoT requires a file in tsv format (Tab-separated values) containing at least the quantified features together with their quantitative values (ID-q.tsv). iSanXoT extracts the required information from the ID-q file to create the relation tables and to integrate the quantified features into higher levels. The majority of proteomics software tools generate tables that can be easily adapted for this purpose (Table 1). The Input Data Adapter module is used to adapt tsv files to create ID-q files that can be directly used by iSanXoT.

**Table 1.**
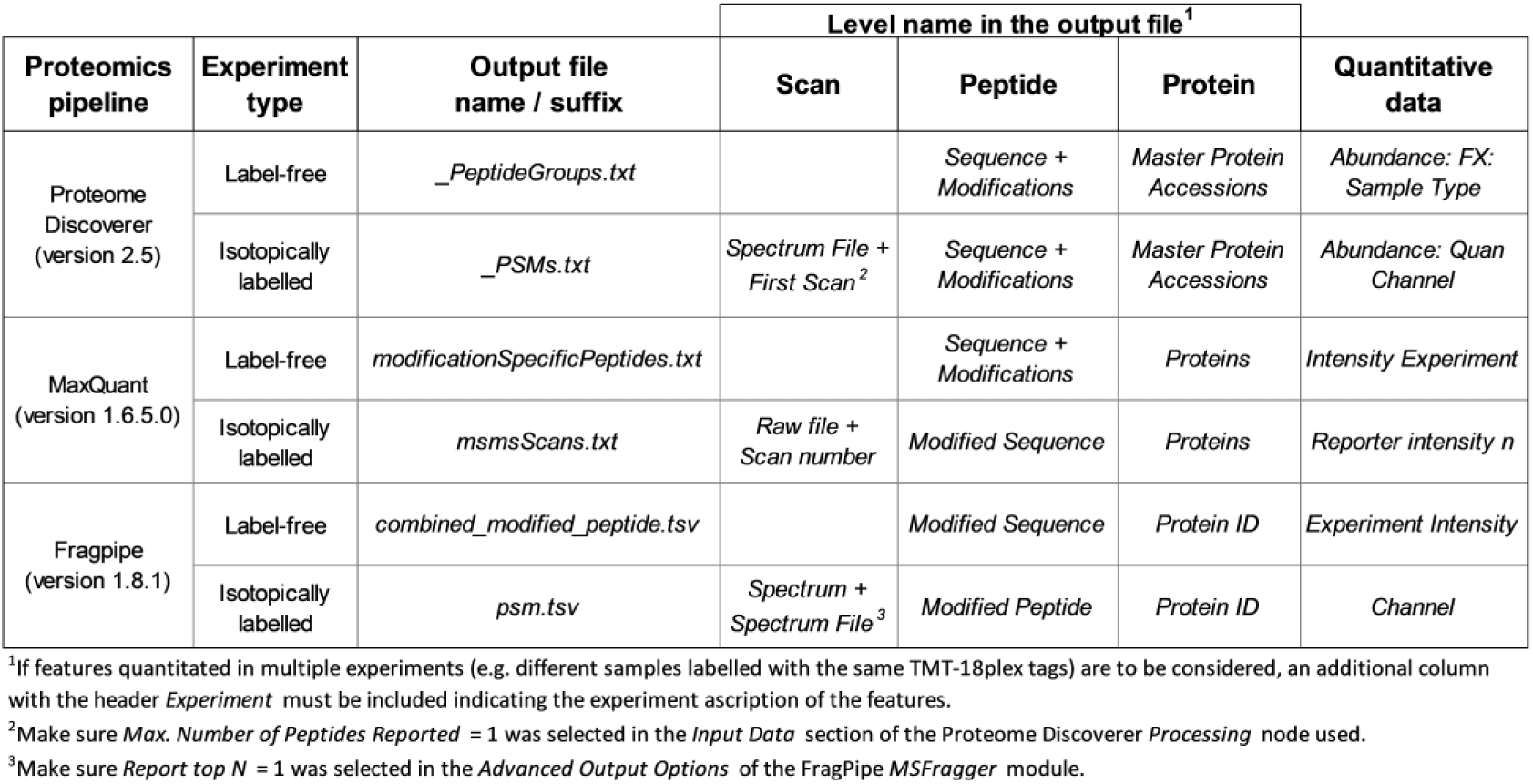
Output data from proteomics pipelines to be included in the ID-q.tsv file.

### Relation Tables Module

This module allows the user to create automatically the relation tables necessary to perform the integrations specified in the *Basic* and *Compound modules* by relating the lower-level elements to the higher-level elements. For instance, the peptide2protein.tsv is the relation table that indicates which peptides belong to each protein. The module displays a task table to be filled in by the user indicating which information from the ID-q file (or other file in tsv format provided by the user) is to be used to create the relation table.

### Basic Modules

The LEVEL CREATOR module is used to define how the ratios between quantitative values are to be calculated from the ID-q table and to assign a name to each ratio.

The LEVEL CALIBRATOR module calibrates the statistical weights (defined as the inverse of variances) associated to each quantitative value at the lowermost level, by fitting a model that describes the relation between signal intensity and variance. The statistical weights are used to control propagation of errors to higher levels and to perform weighted averages at each integration step.

The INTEGRATE module contains a task table where the user indicate which integrations from lower-levels to higher-levels are to be performed using the GIA algorithm [5]. This module also generates a plot per each integration that may be used to check that the distribution of the standardized variables fits the expected normal distribution.

The NORCOMBINE module integrates technical or biological replicates into a single value using the GIA algorithm [5]. Besides calculating weighted averages, GIA integrations take into account error propagation and estimate a global variance across replicates. As above, the accuracy of the integration model can be checked in the normality plots.

The RATIOS module calculates the log2-ratio between the elements of two samples. This step is useful in various contexts to directly compare results obtained between two conditions.

The SBT module applies the Systems Biology Triangle algorithm [5] that performs a triangular integration among a lower level, an intermediate level, and a higher level. The most common application of this module is to apply the SBT model for the detection of changes in functional categories produced by the coordinate behaviour of their proteins. For that, the module uses the protein (lower), category (intermediate) and grand mean (higher) levels.

### Compound Modules

The WSPP-SBT module automatically constructs the relation tables and performs the integrations needed to calculate quantitative protein values using the WSPP model [19] and to apply the SBT algorithm [5]. The WSPP model starts from the quantitative ratios at the scan level, performs the calibration and integrates them to peptide values and then to protein values, and the SBT model is used to detect changes in functional categories due to protein coordination, as depicted above. Since this is one of the most common analysis in quantitative proteomics, we found useful to have a compound module including all the tasks needed.

The three remaining compound modules are extensions of the WSPP-SBT module. The WSPPG-SBT module includes an intermediate protein to gene integration, after which the SBT algorithm is applied at the gene level to unveil coordinated gene behaviour. Analysis at the gene level may have advantages in some situations since functional classification is more elaborated at the gene than at the protein level. In addition, the protein to gene integration avoids dilution of quantitative values due to the presence of isoforms or peptide missassignations and at the same time may detect protein isoforms whose quantitative behavior deviate from the rest of isoforms from the same gene. The WPP-SBT and WPPG-SBT modules are similar to the others, except that the initial calibration is performed at the peptide level. These two last workflows may be useful in label-free experiments quantified at the peptide level.

### Report Modules

The REPORT module generates tsv files displaying the quantitative results at the levels and from the samples selected by the user. This module is highly configurable to include together data from different levels and to allow information from other tsv tables, such as whole protein descriptions or functional classifications, to be included in the reports.

The SANSON module is used to help to resolve functional category redundancy and to visualize global category changes detected by the SBT model. This module generates a similarity graph showing relationships between functional categories on the basis of the protein elements they share

## Results and Discussion

To illustrate iSanXoT performance, we use here three proteomics datasets obtained using isobaric labeling from previous publications in which SanXoT was used for integrative analysis [5, 20, 21], as well as a dataset containing label-free data [24]. We describe below four representative workflows that illustrate the versatility of iSanXoT to perform a variety of quantitative analysis beyond the mere analysis of protein abundances (see the Supplementary Material for a detailed description of these workflows). The workflows and the needed input data from the iSanXoT wiki can be downloaded from GitHub: https://github.com/CNIC-Proteomics/iSanXoT/wiki.

### Workflow 1 (one-step quantification in a labeled experiment)

In this case study, we used quantitative data from García-Marqués *et al*. [5] to illustrate the use of compound modules. This study characterizes the molecular alterations that take place along time when vascular smooth muscle cells (VSMCs) are treated with angiotensin-II (AngII) for 0, 2, 4, 6, 8, and 10 h. Quantitative proteomics was performed using isobaric iTRAQ 8-plex labeling. In this case, we applied the WSPP-SBT module to integrate the quantitative information at the scan, peptide, and protein levels (WSPP model) [6] and to perform a functional category anaysis using the SBT model [5] (Figure 2A). This module works in a fully automated fashion to analyze each one of the samples indicated in the task table. The module calculates relative protein abundance changes (Supplementary Figure S6), producing normally-distributed, standardized quantitative Z values while allowing a full control of error sources at each one of the integration levels (scan, peptide and protein) (see below). The module also applies the Systems Biology Triangle to detect functional protein alterations produced from the coordinated behavior of proteins [5] (Figure 3). Of note, taking advantage of the triangular integration in the SBT model, the module supports improved estimation of protein variance from the protein to category integration [5]. By considering protein variability within functional categories, this method to estimate protein variance is less influenced by the protein changes that take place in the biological model [5].

**Figure 2.**
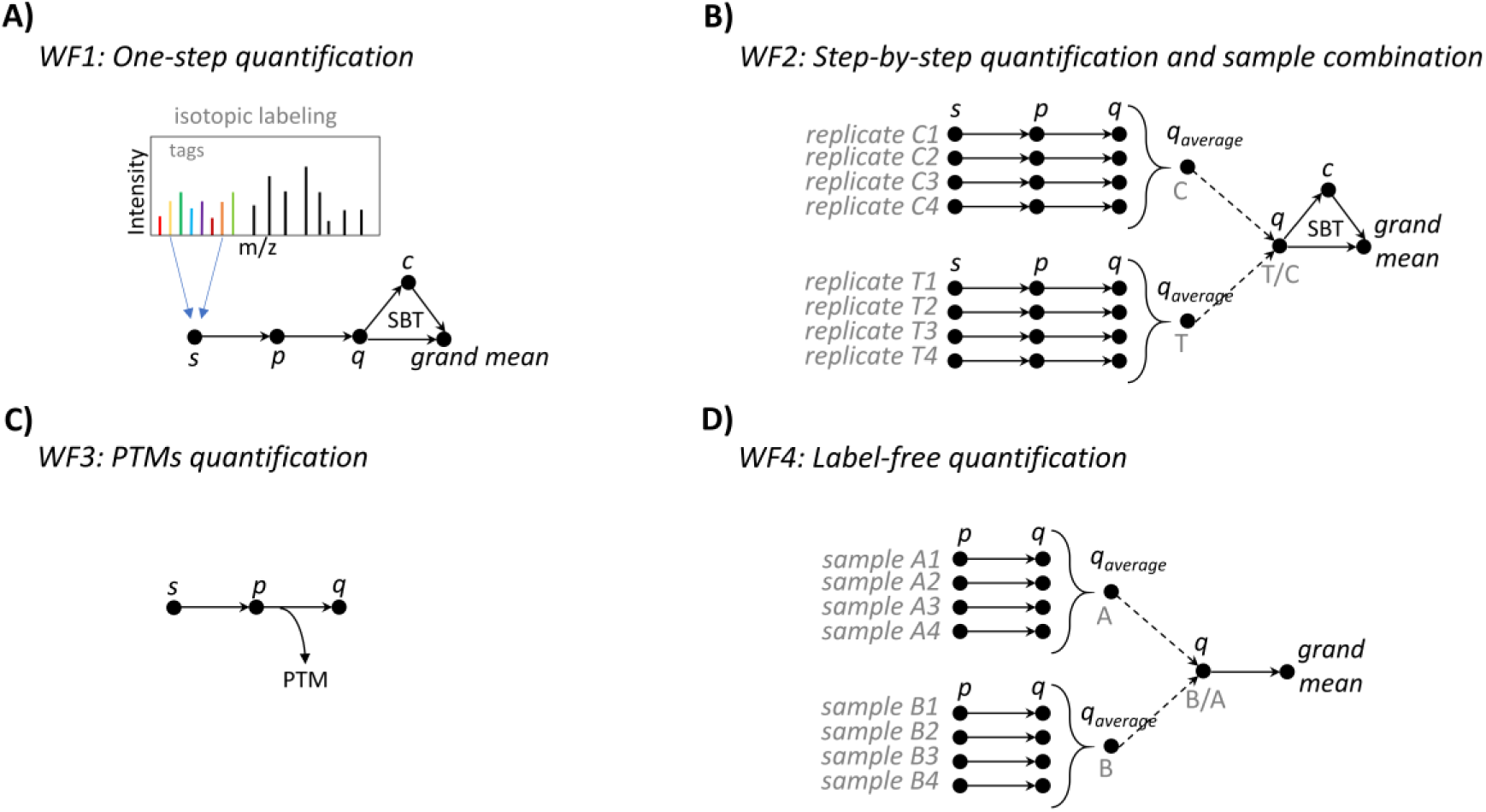
Scheme of the four representative workflows described in this work. These schemes illustrate how the quantitative information is integrated at higher levels or combined across sample types in each workflow. Note that while WF1 (A) is performed using a compound module that automatically performs all required steps, the other three workflows (B-D) were constructed by indicating step-by-step the tasks to be performed through the interface. The rationale of these workflows is described in Results and Discussion. More specific details about how to use the interface are provided in the Supplementary Information file. All these workflows can be reused or adapted to other sample configurations according to user needs.

**Figure 3.**
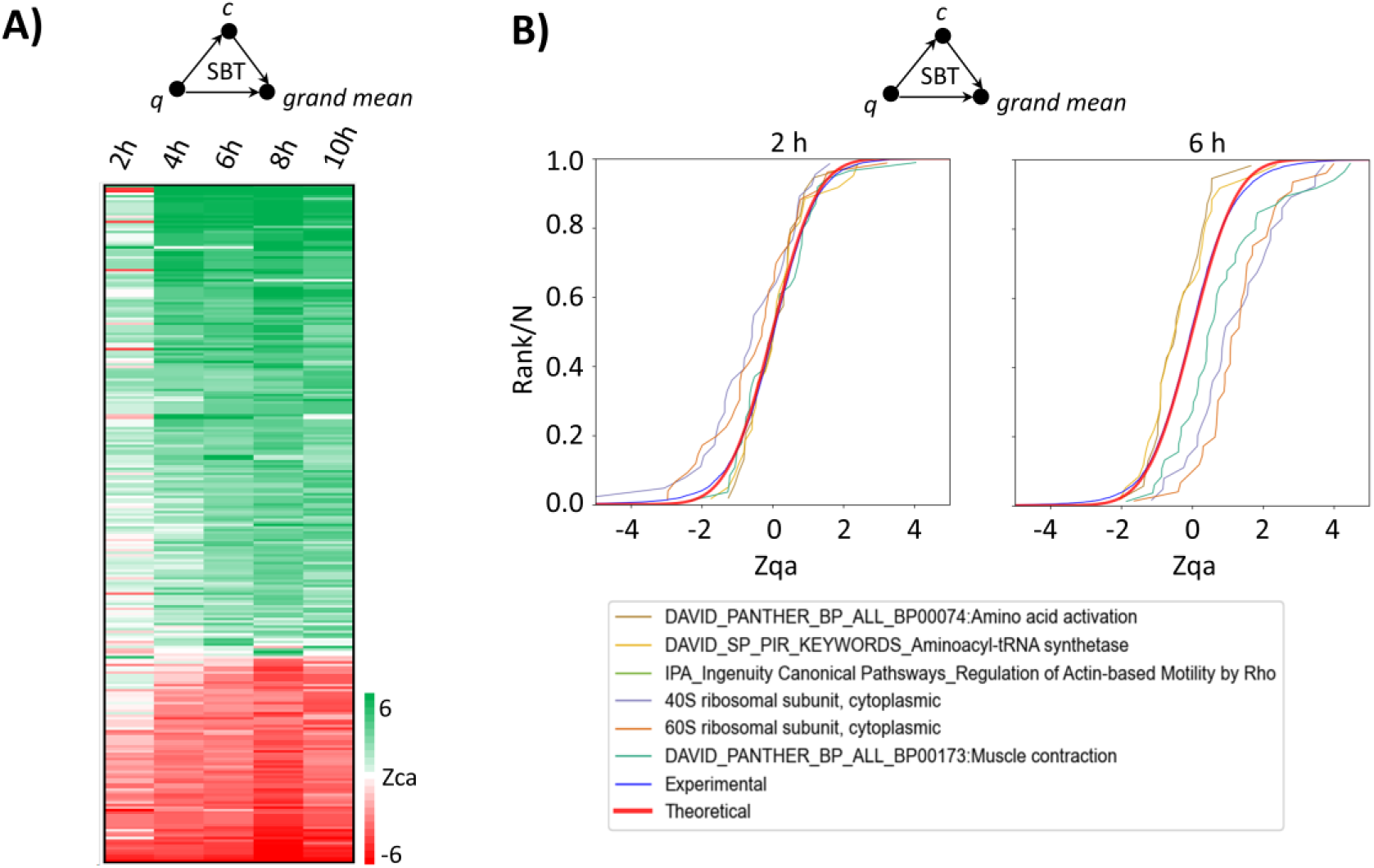
Representative results of workflow 1 obtained using iSanXoT. A) Left panel: Heatmap depicting the changes in functional categories produced by the coordinated behaviour in proteins along time. B) Examples of functional category changes in relation to the null hypothesis at 2h and 6h after incubation with angiotensin-II. A and B are results obtained by applying workflow 1.

### Workflow 2 (step-by-step quantification and sample combination in a labeled experiment)

In this application, we used the data from the study performed by González-Amor *et al*. study[21], to illustrate how the full list of *Basic modules* can be used to construct step-by-step a quantitative workflow. In this case the workflow is designed to calculate protein values in several samples by integrating in each one the quantitative data at the scan, peptide and protein levels (Figure 2B). The workflow then integrates the quantitative information from technical or biological replicates within user-defined sample groups, and analyzes protein abundance changes across these groups (Figure 2B). Finally, the workflow captures category abundance changes produced by coordinated behaviour of proteins (Figure 2B). As previously shown [3], the workflow revealed a coordinated alteration of proteins implicated in cardiovascular function, extracellular matrix and remodeling, and vascular redox state in aortic tissue from AngII-infused ISG15-KO mice (Figure S20A). The coordinated protein behavior from some of the altered categories can be easily analyzed in the sigmoid plots (Figure S20B).

This workflow also illustrates two relevant features of the statistical approach followed by iSanXoT. The first one is the automated calibration step (performed by the LEVEL CALIBRATOR module), where a statistical weight is assigned to each quantitative measure at the initial (scan) level. The statistical weight is the inverse of the variance of each quantification and is estimated from the signal intensity by fitting a global model. iSanXoT automatically generates plots which allow testing the accuracy of the model (Supplementary Figure S12). This initial callibration step allows subsequent integrations to be performed using weighted averages and controlling error sources at all the levels. The second one is the ability of iSanXoT to integrate protein values from several samples (NORCOMBINE module) applying the GIA algorithm, which performs a weighted averaging and accurately models the dispersion of protein values around the average taking into account error propagation and allowing a full control over outliers [19] (Figure 4). This unique design allows the integration of protein values obtained in non-balanced samples, different experiments or mass spectrometers and even different labelling techniques, as we showed previously [19].

**Figure 4.**
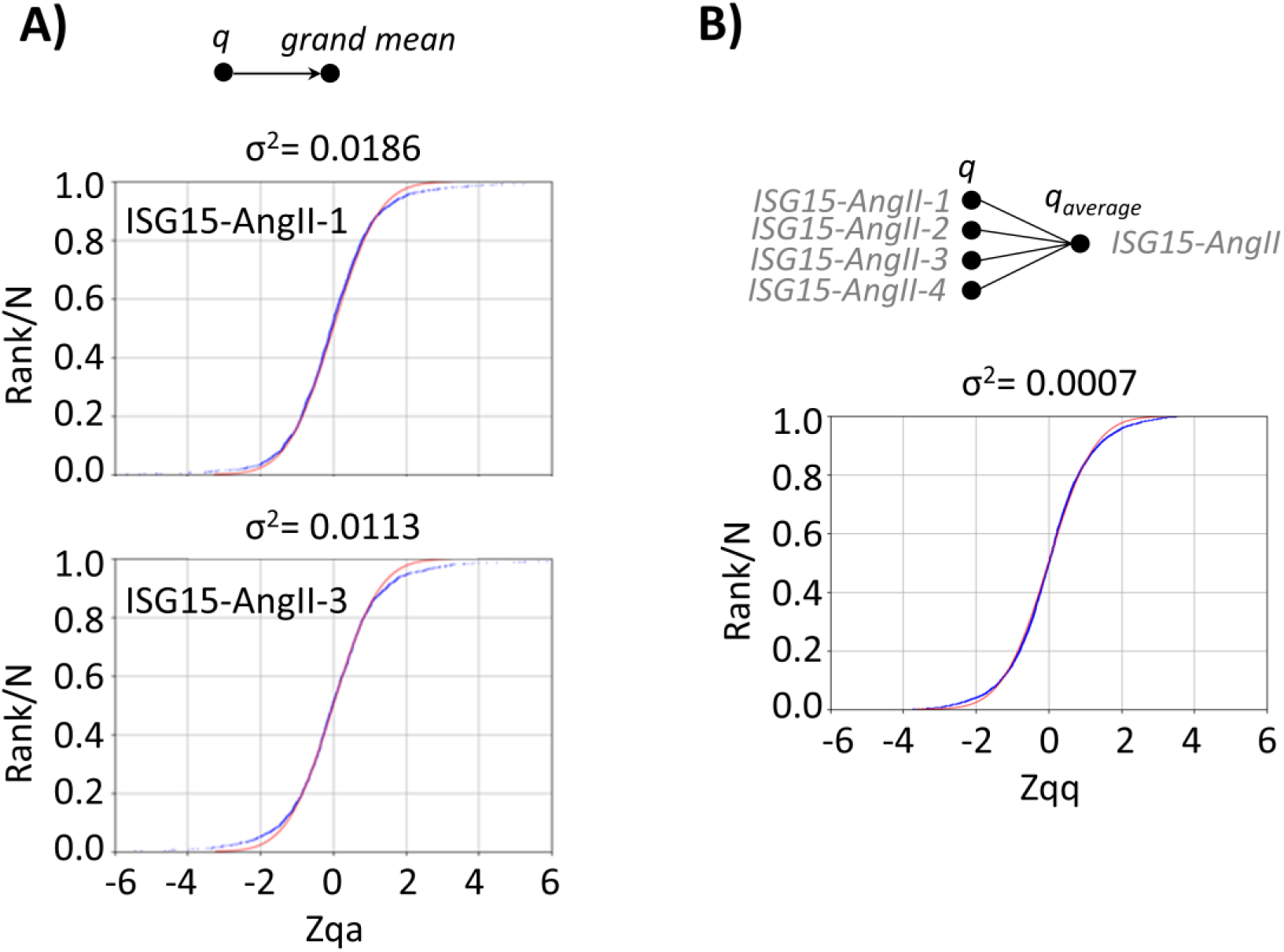
Representative results of workflow 2. A) Example of the distribution of standardized protein abundance values showing the good alignment with the theoretical prediction of the model taken from sample ISG15-AngII-4 from workflow 2. B) Distribution of the standardized variable at protein level around the averages in the process of integration of replicates from the sample group ISG15-AngII, showing the good accuracy of the GIA algorithm.

### Workflow 3 (quantification of posttranslationally modified peptides in a labeled experiment)

This workflow is designed to analyze quantitative data of posttranslational modifications obtained by Bonzon-Kulichenko *et al*. [20]. In this study FASILOX, a redox proteomics technique was used to analyze reversible oxidation on Cys residues in mouse embryonic fibroblast preparations subjected to chemical oxidation. FASILOX labels reduced and oxidized Cys residues with different alkylating reagents, allowing these two group of Cys sites to be quantified separately [20]. This workflow makes the conventional integrations from scan to peptide and from peptide to protein levels, and concentrates on the results of the GIA algorithm to model the dispersion of peptide values when they are integrated to their proteins, in order to detect peptidoforms that significantly deviate from the expected behavior, independently from protein abundance changes (Figure 2C). Of note, the standardized variable that the model uses to describe the quantitative behaviour of all the peptidoforms in the experiment follows very accurately the expected normal distribution (Figure 5A, red and blue lines). Hence, this workflow is very useful to detect statistically significant abundance changes of post-translationally modified peptides, decoupling them from the proteins they come from, on the basis of a robust and validated statistical framework [7, 8] (Figure 5A, brown and green lines). For simplicity we show here a simple application to redox proteomics [7], but the same workflow can also be used for high-throughput PTM data obtained, for instance, from open searches, as we have demonstrated previously [8].

**Figure 5.**
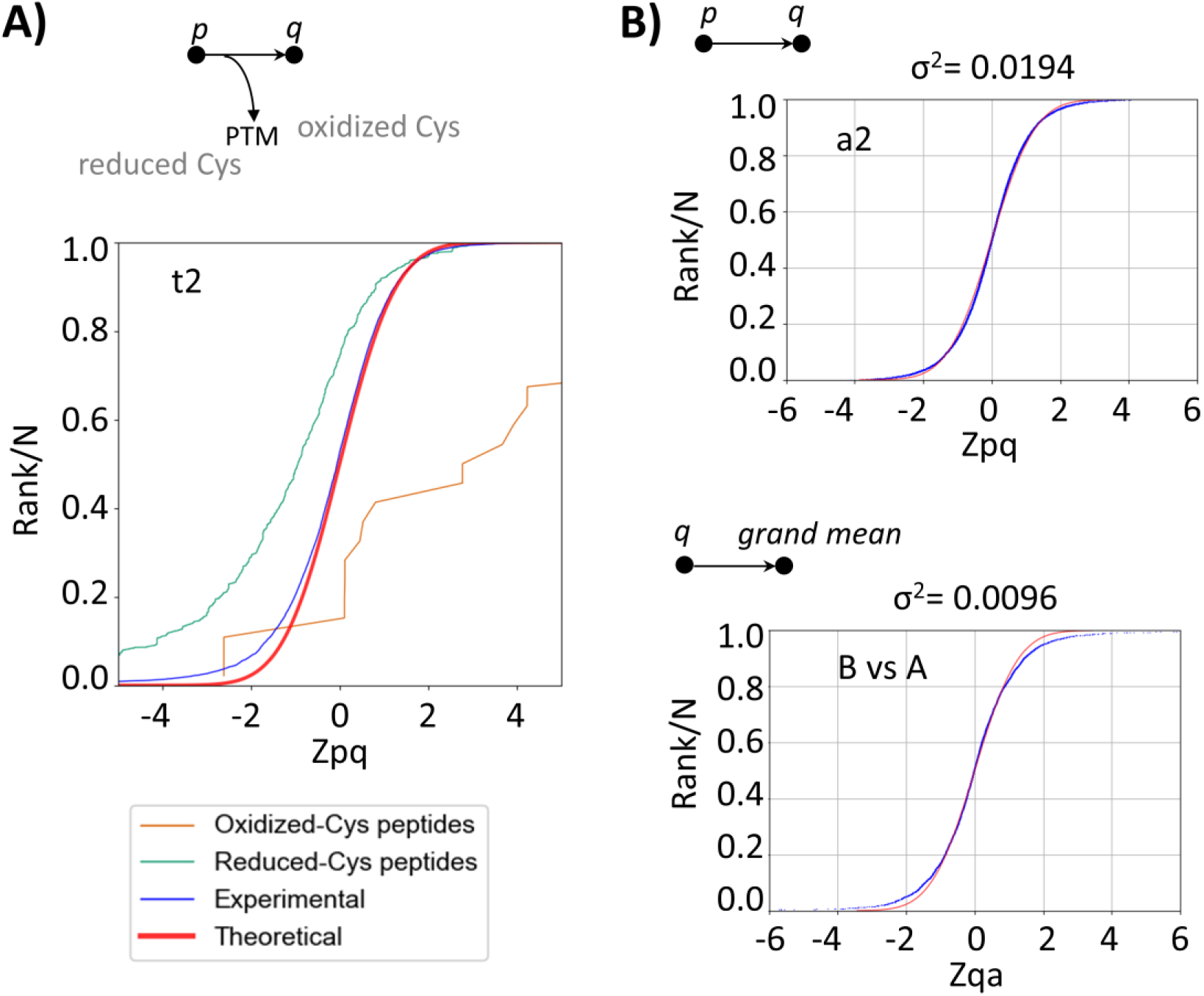
Representative results of workflow 3 and workflow 4. A) Distribution of the standardized variable at the peptide level around the protein averages for all the peptidoforms of the proteome (blue), of oxidized Cys-containing peptides (brown) and of reduced-Cys containing peptides (green) in one sample of the FASILOX experiment (workflow 3). Note that the model captures with very good accuracy the distribution of peptide quantifications and how the oxidized and reduced forms tend to increase or decrease, respectively, deviating from the expected theoretical distribution (red). B) Distribution of the standardized variable at the peptide (top) and protein levels (bottom) from one of the samples in workflow 4, showing the good accuracy of the model to describe quantitative variability in label-free experiments.

### Workflow 4 (label-free quantification)

In this application, we used data from Navarro *et al*. [24], to illustrate the application of iSanXoT to analyze quantitative data obtained using label-free approaches. In this study two mixtures of human, yeast and Escherichia coli proteins at different proportions were compared. The MS data was processed to peptide level using MaxQuant. The output from MaxQuant was adapted to be used by iSanXoT using the Input Data Adapter module. The iSanXoT worflow was designed to analyze protein abundance changes between the two sample groups by transforming the quantitative peptide data into relative abundances at the peptide level and then performing subsequent integrations as in workflow 2 (Figure 2D). This is an alternative approach to analyze label-free data that has advantages in some situations. Modeling of variances, normalization, standardization and statistical weighting are performed automatically, without data filtering, pre-processing or value imputation, using the GIA algorithm (Figure 5B, top), even in situations of highly imbalanced data. As above, integration of data from different experiments is also possible (Figure 5B, bottom). Finally, iSanXoT also supports selective normalization on the basis of a subset of data, such as a group of proteins of the same procedence (human in Supplementary Figure S32).

## Conclusions

iSanXoT exploits the unique capabilities of the GIA algorithm for statistical analysis of quantitative proteomics data, extending the range of analytical resources of conventional quantitative packages. Since each GIA integration performs an independent statistical modeling and standardization of the integrated data, the quantitative information can be limitlessly extended to superior levels or integrated across samples within a given level, maintaining a full control of error sources at each step. By performing serial GIA integrations, this approach naturally allows for the creation of quantitative workflows adaptable to a large variety of user’s needs.

In this work we have selected four experiments that illustrate how iSanXoT can be used to create different workflows designed to answer specific biological questions. These include the automated analysis of functional categories that change as a result of the coordinated action of proteins. This kind of analysis require a proper statistical model that captures the underlying protein variance within the functional categories used to classify the proteins and that is automatically applied in iSanXoT by a dedicated module. In other of the examples presented we illustrate the application of the GIA algorithm to integrate and compare protein values across different sample types. The application of the GIA algorithm for the analysis of PTM is presented in another example, where we demonstrate the good accuracy of the statistical model to describe the quantitative variance at the peptide level. The high-throughput quantitative analysis of post-translational modifications is a field that is attracting considerable interest, particularly with the development of open-search approaches. Finally, an application to the analysis of label-free data is also presented; this application may be used as an alternative to conventional approaches with some advantages in some cases, for instance when the data is highly balanced or when a careful control of variance at the peptide level is needed.

iSanXoT is a standalone application with an user-friendly interface, enabling versatile and highly automatable creation of integrative quantification workflows and of report tables. Once prepared, the workflows can be exported and straightforwardly reused across different projects. The current version of the software accepts the direct output from several widespread proteomics pipelines, making it readily available for the proteomics community.

## Authors’ contributions

JMR was responsible for conceptualizing the project, developing the methodology and software, curating the data, conducting formal analysis, validating the results, writing the original draft, and performing review and editing tasks. EC conducted validation, managed data curation, and contributed to the review and editing processes. IJ handled validation and participated in editing. RBR and RM developed certain programs. EN, AL, CAD, and JAL were involved in validation and data curation efforts. JV played a role in conceptualization, acquired funding, provided supervision, contributed to review writing, and participated in editing.

## Acknowledgements

This study was supported by competitive grants from the Spanish Ministry of Science, Innovation and Universities (PGC2018-097019-B-I00, PID2021-122348NB-I00, PLEC2022-009235 and PLEC2022-009298), the Instituto de Salud Carlos III (Fondo de Investigación Sanitaria grant PRB3 (PT17/0019/0003-ISCIII-SGEFI / ERDF, ProteoRed), Comunidad de Madrid (IMMUNO-VAR, P2022/BMD-7333) and “la Caixa” Banking Foundation (project codes HR17-00247 and HR22-00253). The CNIC is supported by the Instituto de Salud Carlos III (ISCIII), the Ministerio de Ciencia e Innovación (MCIN) and the Pro CNIC Foundation), and is a Severo Ochoa Center of Excellence (grant CEX2020-001041-S funded by MICIN/AEI/10.13039/501100011033).

