## Supplementary Material for "iSanXoT: a Standalone Application for the Integrative Analysis of Mass Spectrometry-Based Quantitative Proteomics Data"

---

### Sample Workflows with Application to Case Studies

We describe below in detail four sample workflows that illustrate the capacity of iSanXoT to statistically ascertain protein or peptide abundance changes in a variety of biological contexts. Note that these workflows may be easily reused to process new data (see next section).

#### Workflow 1: One-step quantification in a labeled experiment

##### Experimental

The identification and quantification data from García-Marqués *et al*. [1] were used to illustrate this workflow. This study characterizes the molecular alterations that take place along time when vascular smooth muscle cells (VSMCs) are treated with angiotensin-II (AngII) for 0, 2, 4, 6, 8, and 10 h. Quantitative proteomics was performed using isobaric iTRAQ 8-plex labeling. Workflow 1 analyzes a) protein abundance changes and b) functional category alterations produced by the coordinated behaviour of proteins at each one of the times, in relation to time 0. This is done using, in only one step, the compound module WSPP-SBT, which performs automatically all the required tasks.

##### Workflow operation

Workflow 1 requires the RELS CREATOR module, the WSPP-SBT compound module and the REPORT basic module (*Figure S1*). The relation tables required to perform the integrations are created by the RELS CREATOR module (*Figure S1*A) from a table provided by the user. The WSPP-SBT module performs a sequence of consecutive integrations based on the WSPP statistical model [2] and the SBT algorithm [1] (*Figure S1*B). Finally, the REPORT module organizes the data in tables containing the information needed.

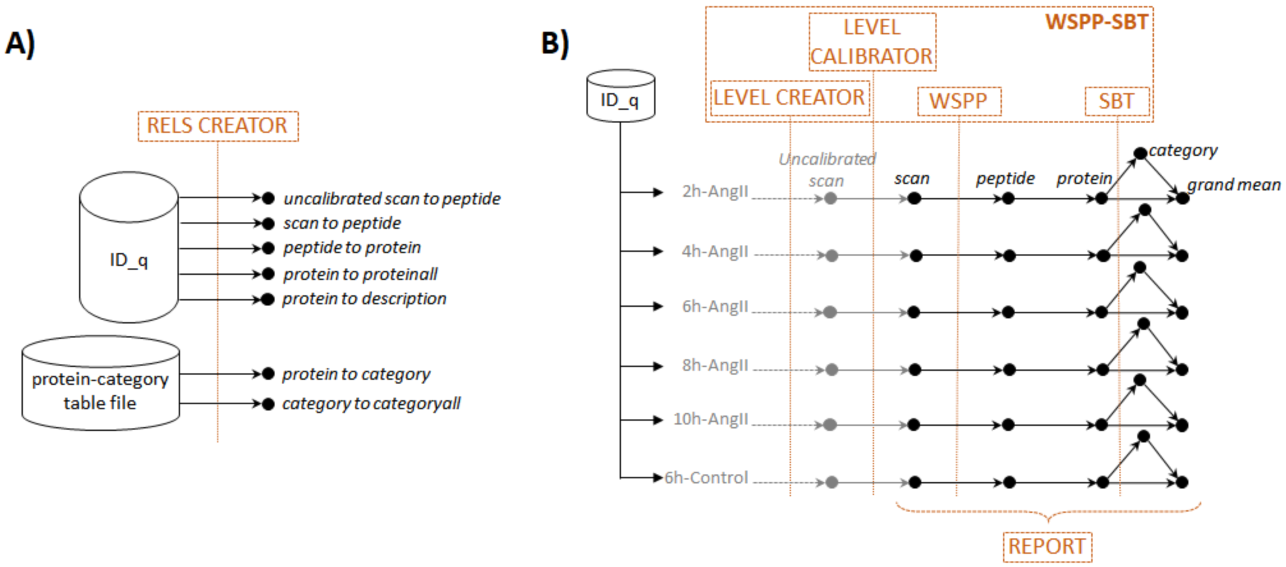

*Figure S1. Scheme of workflow 1 (one-step quantification in a labeled experiment) showing module components: RELS CREATOR (A) and WSPP-SBT and REPORT (B)*

The WSPP-SBT module needs the user to define the meaning of relative abundances, which iSanXoT always expresses as log2ratios. In this case the abundance data corresponds to the intensities of iTRAQ reporters at the scan level, which are tabulated in the "ID-q" file with the name of each reporter as column header (see below how these tables are generated). The intensities of each scan at 0 h are in column "Abundance: 113" and are used as a common reference to express abundance ratios and are therefore used as denominator. The reporter intensities corresponding to the different time points are used as numerators of the ratios. The task table also allows the user to put an easily identifiable name in the folders where the quantitative values of each sample are stored (*Figure S2*).

| Experiment | Identifier column header | Ratio numerator | Ratio denominator | Output Sample folder |
|---|---|---|---|---|
| VSMC | Scan_Id | Abundance: 114 | Abundance: 113 | 2h-AngII |
| VSMC | Scan_Id | Abundance: 115 | Abundance: 113 | 4h-AngII |
| VSMC | Scan_Id | Abundance: 116 | Abundance: 113 | 6h-AngII |
| VSMC | Scan_Id | Abundance: 117 | Abundance: 113 | 6h-Control |
| VSMC | Scan_Id | Abundance: 118 | Abundance: 113 | 8h-AngII |
| VSMC | Scan_Id | Abundance: 119 | Abundance: 113 | 10h-AngII |

*Figure S2. The WSPP-SBT task table for workflow 1.*

The WSPP-SBT module first performs a calibration to assign a statistical weight to each one of the log2ratio values at the scan level (*Figure S1*B), as described [2]. The statistical weight of each scan is the inverse of the estimated variance associated with the log2 of intensity ratios [2]. Once the data is calibrated at the scan level, the workflow performs the integrations *scan-to-peptide* and *peptide-to-protein*.

At the protein level the SBT algorithm is then applied for the detection of functional category changes originated by the coordinated behaviour of proteins (*Figure S1*B). The algorithm first calculates the variance of the *protein-to-category* integration, which is an improved estimate of the technical protein variance, since it is less influenced by biological changes [1]. This protein variance is used to perform the *protein-to-grand mean* integration (hereinafter referred to as *protein-to-proteinall*) integration, from which statistically significant abundance changes are detected. The algorithm finally performs the *category-to-grand mean* integration (hereinafter referred to as *category-to-categoryall*), from which statistically significant category changes are detected. All the results from the integrations executed by the WSPP-SBT module are saved, per each sample, to the *Output Sample folder* indicated in the module task table (*Figure S2*).

Every integration step needs a relation table (a text file) that links lower- to higher-level elements. Relation tables can be automatically created by the RELS CREATOR module (*Figure S1*A, upper) and can be provided by the user (*Figure S1*A, lower). In this example (*Figure S3*) the relation tables linking scan to peptides and peptides to proteins are obtained from the "ID-q" file, just by indicating the name of the columns where they are located (in this case "Scan_Id", "Pep_Id" and "Master Protein Accessions"). In this case the columns *Master Protein Accessions* and *Master Protein Descriptions* in the "ID-q" file contain the accession numbers and the complete name of the proteins, respectively, so that a relation table *protein2description* is also created; this relation table may be later used to append the full name of the protein to any of the created reports (see below). An example of the *peptide2protein* relation table, linking the peptides identified to the proteins they come from is showed in *Figure S4*A. The elements of the relation table protein2category were retrieved from a text file containing functional annotations for mouse proteins gathered from several protein function databases (*Figure S4*B), as described by the authors [1]. Note that relation tables are by default extracted from the ID-q file; to use other text files the absolute path with the location of the text file has to be indicated. The relation tables *protein2proteinall* and *category2categoryall* guide the integration to a grand mean (a common element called "[1]"). The integration *peptide2peptideall* is not necessary in this workflow but is included in this example since it may be useful to inspect quantifications at peptide level.

| Relation Table to be created | Column name of Lower level | Column name of Higher level | Column name of 3rd c | Table from which RT is extracted |
|---|---|---|---|---|
| uscan2peptide | Scan_Id | Pep_Id | | |
| scan2peptide | Scan_Id | Pep_Id | | |
| peptide2protein | Pep_Id | Master Protein Accessions | | |
| peptide2peptideall | Pep_Id | [1] | | |
| protein2proteinall | Master Protein Accessions | [1] | | |
| protein2description | Master Protein Accessions | Master Protein Descriptions | | |
| protein2category | Protein | Category | | {PATH}/DAVID_IPA_merged_noDups_feb14_IJ.txt |
| category2categoryall | Category | [1] | | {PATH}/DAVID_IPA_merged_noDups_feb14_IJ.txt |

*Figure S3. The RELS CREATOR task table for workflow 1.*

A)

| protein | peptide |
|---|---|
| P23242 | KVAAGHELQPLALVDQRPSSR__N-Term(iTRAQ8plex);K1(iTRAQ8plex) |
| P17182 | AAVPSGASTGLYEALELRDNDKTR__N-Term(iTRAQ8plex);K22(iTRAQ8plex) |
| P23780 | AGATLDLLVENMGR__N-Term(iTRAQ8plex) |
| A2AIM4; P58771-2; P58774-2; E9Q45 | RLQLVEEELDRAQER__N-Term(iTRAQ8plex) |
| P20152 | EKLQEEMLQREEAESTLQSFR__N-Term(iTRAQ8plex);K2(iTRAQ8plex) |
| P99024 | MAVTFLGNSTALQELFKR__N-Term(iTRAQ8plex);M1(Oxidation);K17(iTRAQ8plex) |
| A1BN54; Q9JI91; P57780; Q7TPR4 | KHEAFESDLAAHQDR__N-Term(iTRAQ8plex);K1(iTRAQ8plex) |
| Q91YQ5 | ASSFVLALEPELESR__N-Term(iTRAQ8plex) |
| O08547 | NLGSLNTELQDVQR__N-Term(iTRAQ8plex) |
| Q9EQ06 | SVAGELVLLTGAGHGLGR__N-Term(iTRAQ8plex) |
| P05064; Q9CPQ9 | FSNEELAMATVTALR__N-Term(iTRAQ8plex) |
| P10126 | KDGSASGTTLLEALDCLLPPTRPTDKPLR__N-Term(iTRAQ8plex);C16(Carbamidomethyl);K26(iTRAQ8plex) |

B)

| category | protein |
|---|---|
| DAVID_PANTHER_BP_ALL_BP00141:Transport | Q9Z351 |
| DAVID_PANTHER_BP_ALL_BP00142:Ion transport | Q9Z351 |
| DAVID_PANTHER_BP_ALL_BP00143:Cation transport | Q9Z351 |
| DAVID_PANTHER_PATHWAY_P00042:Muscarinic acetylcholine receptor 1 and 3 s | Q9Z351 |
| DAVID_SP_PIR_KEYWORDS_alternative splicing | Q9Z351 |
| DAVID_SP_PIR_KEYWORDS_ion transport | Q9Z351 |
| DAVID_SP_PIR_KEYWORDS_ionic channel | Q9Z351 |
| DAVID_KEGG_PATHWAY_mmu04020:Calcium signaling pathway | Q9Z329 |
| DAVID_KEGG_PATHWAY_mmu04070:Phosphatidylinositol signaling system | Q9Z329 |
| DAVID_KEGG_PATHWAY_mmu04114:Oocyte meiosis | Q9Z329 |
| DAVID_KEGG_PATHWAY_mmu04270:Vascular smooth muscle contraction | Q9Z329 |

*Figure S4. Excerpt from the peptide2protein (A) and protein2category (B) relation tables that link peptides to proteins and proteins to categories, respectively.*

Once the integrations are performed, the REPORT module is used to collect from the *Output sample folders* stated by the user the statistical variables desired and to organize them in tables (*Figure S5*). In this case, the results from the samples (2h-AngII, 4h-AngII, 6h-AngII, 8h-AngII, and 10h-AngII) are to be tabulated.

In this example the REPORT module creates a protein table and a category table by performing the following steps:

- Create a table called “Npep2prot" containing the number of peptides with which each protein is quantified.
  - This is done by extracting from the *peptide-to-protein* integrations in the indicated folders the number of elements (n) of the lower level (peptide) used to quantitate the higher level (protein).
- Create a table called “Npep2prot_Quantprot_filtered” containing the protein changes Zqa and the statistical significance FDRqa of these changes.
  - This is done by extracting from the *protein-to-proteinall* integration in the indicated folders the standardized log2 ratios (Z) and False Discovery Rates (FDR) of the lower level (protein).
- Add to this table the number of peptides with which each protein is quantified.
  - This is done by merging the previous table with the existing table “Npep2prot” according to the level common to the two tables (protein), without including a specific column (peptide) and eliminating replicate entries.
- Add to this table an additional column with the complete description of the proteins.

    - This is done by merging the previous table with the relation table protein2description according to the level common to the two tables (protein).
- Filter the table so that only the proteins having a statistically significant abundance change (FDR < 0.01) are tabulated.
    - This is done by applying in the Filter column a condition based on the FDR to the results from the protein2proteinall integration. For more detailed information, see the “Filter for report” in the iSanXoT wiki: https://github.com/CNIC-Proteomics/iSanXoT/wiki.

- Create a table called “Nprot2cat” containing the number of proteins with which each category is quantified.
    - This is done by extracting from the *protein-to-category* integrations in the indicated folders the number of elements (n) of the lower level (protein) used to quantitate the higher level (category).

- Create a table called “Nprot2cat_Quantcat_filtered” containing the category changes Zca and the statistical significance FDRca of these changes.
    - This is done by extracting from the *category-to-categoryall* integration in the indicated folders the standardized log2 ratios (Z) and False Discovery Rates (FDR) of the lower level (category).
- Add to this table the number of proteins with which each category is quantified.
    - This is done by merging the previous table with the existing table “Nprot2cat” according to the level common to the two tables (category), without including a specific column (protein) and eliminating replicate entries.
- Filter the table so that only the categories having a statistically significant change (FDR < 0.01) are tabulated.
    - This is done by applying in the Filter column a condition based on the FDR to the results from the category2categoryall integration.

- Create a table called “Npep2prot_Quanprot” containing the number of peptides per protein, the protein changes Zqa and the statistical significance FDRqa of these changes.
    - This is done as explained above, omitting the protein descriptions and the filters.

- Create a table called “Nprot2cat_Quancat_Quanprot_filtered” containing the category changes Zca and the statistical significance FDRca of these changes.
    - This is done by extracting from the *category-to-categoryall* integration in the indicated folders the standardized log2 ratios (Z) and False Discovery Rates (FDR) of the lower level (category).
- Add to this table the number of proteins per category, the protein changes Zqa and the statistical significance FDRqa of these changes.
    - This is done by merging the previous table with the existing tables “Nprot2cat” and “Npep2prot_Quantprot”.
- Filter the table so that only the categories containing 5 or more proteins or 100 or less proteins are tabulated.
    - This is done by applying in the Filter column a set of conditions joined with the “&” operator.

Note that these commands in the REPORT module, which allow to construct tables required in typical quantitative proteomics projects, are easily reusable for other projects.

| Sample folder(s) | Lower level | Higher level | Reported vars | Output report | Column headers to eliminate | Merge with report | Add columns from relation tal | Filter |
|---|---|---|---|---|---|---|---|---|
| 2h-AngII , 4h-AngII , 6h-AngII , 8h-AngII , 10h-AngII | peptide | protein | n | Npep2prot | | | | |
| 2h-AngII , 4h-AngII , 6h-AngII , 8h-AngII , 10h-AngII | protein | proteinall | Z , FDR | Npep2prot_Quanprot_filtered | peptide | Npep2prot | protein2description | FDR_protein2proteinall < 0.01 |
| 2h-AngII , 4h-AngII , 6h-AngII , 8h-AngII , 10h-AngII | protein | category | n | Nprot2cat | | | | |
| 2h-AngII , 4h-AngII , 6h-AngII , 8h-AngII , 10h-AngII | category | categoryall | Z , FDR | Nprot2cat_Quancat_filtered | protein | Nprot2cat | | FDR_category2categoryall < 0.01 |
| 2h-AngII , 4h-AngII , 6h-AngII , 8h-AngII , 10h-AngII | protein | proteinall | Z , FDR | Npep2prot_Quanprot | peptide | Npep2prot | | |
| 2h-AngII , 4h-AngII , 6h-AngII , 8h-AngII , 10h-AngII | category | categoryall | Z , FDR | Nprot2cat_Quancat_Quanprot_filtered | | Nprot2cat , Npep2prot_Quanprot | | (n_protein2category >=5) & (n_protein2category <= 100) |

*Figure S5. The REPORT task table for workflow 1.*

In *Figure S7* we show two heat maps constructed from the protein and category tables obtained with the REPORT module.

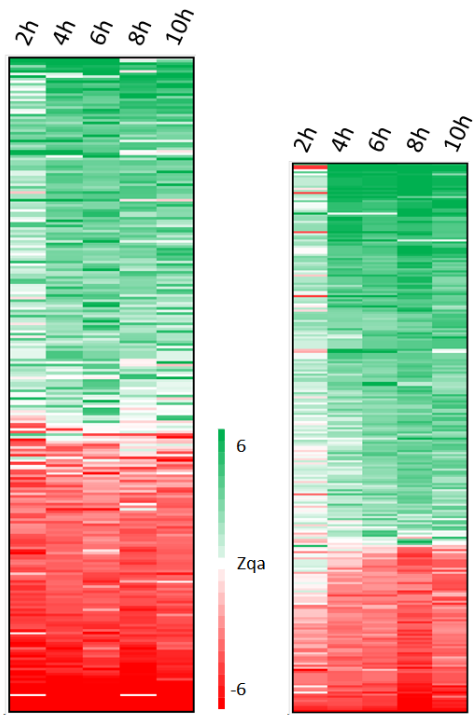

*Figure S6. Relative abundance changes of proteins (Zqa, Left) and functional categories (Zca, Right) generated from the "Npep2prot_Quanprot_filtered" and "Nprot2cat_Quancat_filtered" reports, respectively, obtained by the REPORT module of workflow 1 (Figure S5). Both report tables were sorted by the average of Zqa and Zca, respectively.*

In Figure S7 we show some examples of functional categories showing statistically significant changes produced by coordinated protein behaviour, plotted using the data in the “Nprot2cat_Quancat_Quanprot_filtered" table generated by the REPORT module.

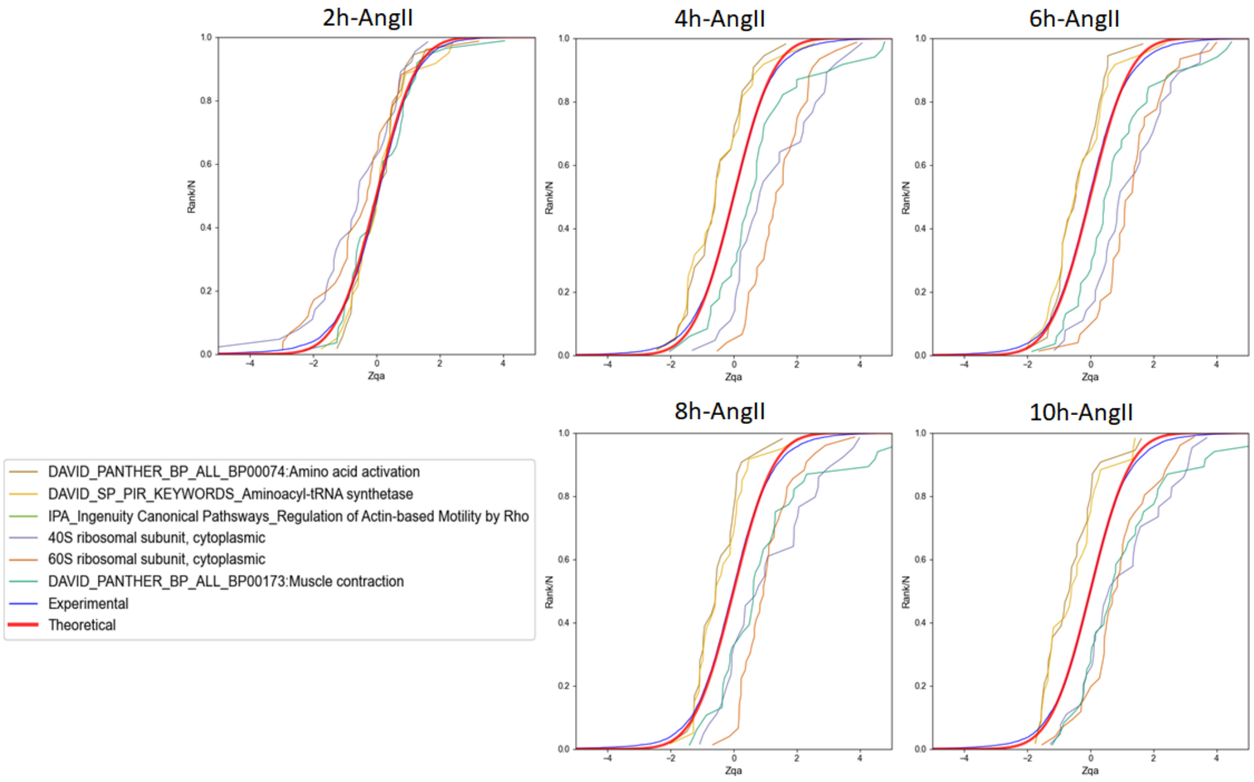

*Figure S7. Examples of time-dependent coordinated protein behavior of VSMCs treated with angiotensin-II as revealed by the distribution of the standardized log2 ratio (Zqa) of protein components in each category.*

#### Workflow execution

The workflow template and input files that are needed to execute this workflow can be downloaded from

https://github.com/CNIC-Proteomics/iSanXoT/wiki/studies/cases/templates/WSPP-SBT.zip. See the *Importing a workflow template* Section below for detailed instructions.

### Workflow 2: Step-by-step quantification and sample combination in a labeled experiment

#### Experimental

The data from González-Amor *et al*. [3] was used to illustrate this workflow. This study analyzes the contribution of interferon-stimulated gene 15 (ISG15) to the vascular damage associated with hypertension, using KO mutants for this gene and subjecting or not the animals to AngII treatment. This experiment contained 16 samples from mouse aortic tissue corresponding to four groups: four WT-Control mice, four ISG15-KO mice, four WT+AngII mice, and four ISG15-KO+AngII mice. The experiment was performed using two isobaric iTRAQ 8-plex batches. This module illustrates how to create, step by step, a workflow to integrate the quantitative results from each sample to the protein level, to integrate protein data from the four biological replicates in each group, to construct ratios between two conditions, and to analyze functional category changes due to coordinated protein behavior using the SBT model.

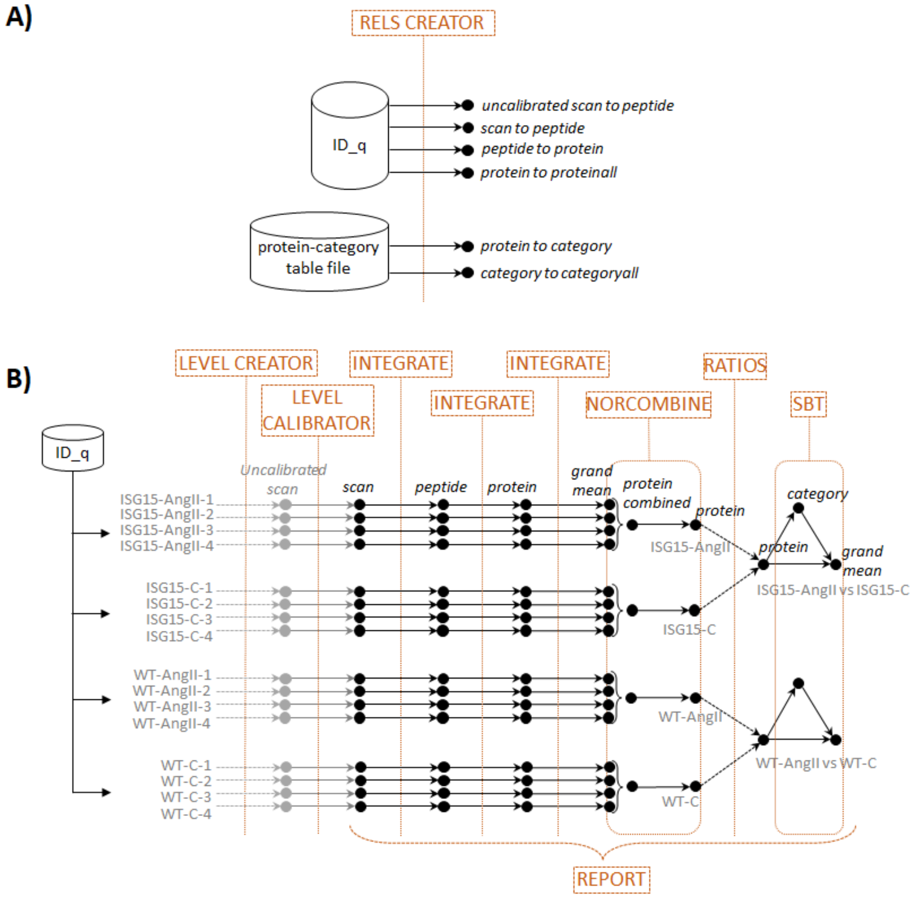

*Figure S8. Scheme of workflow 2 (step-by-step quantification and sample combination in a labeled experiment) showing module components: RELS CREATOR (A) and LEVEL CREATOR, LEVEL CALIBRATOR, INTEGRATE, NORCOMBINE, RATIOS, SBT, and REPORT (B)*

#### Workflow operation

Workflow 2 comprises all six basic modules: LEVEL CREATOR, LEVEL CALIBRATOR, INTEGRATE, NORCOMBINE, RATIOS, and SBT as well as the REPORT module (*Figure S8*). The task table in the starting module, LEVEL CREATOR, generates the files at the scan level containing the log2 ratios and the corresponding sample folders (*Figure S9*). In this example, as in workflow 1, the name of each iTRAQ reporter was used as column header in the “ID-q” file containing the intensities. In addition, the column Experiment indicates whether the intensities come from the first or second iTRAQ 8-plex batch. Each iTRAQ batch contains two biological replicates from each of the four groups and the average of reporter intensities from the two untreated WT mice (reporters in the columns “113” and “117”) were used as internal control within each batch, and therefore were used as denominator for the log2ratios.

| Experiment | Identifier column header | Ratio numerator column | Ratio denominator column(s) | Level to be created | Output Sample folder |
|---|---|---|---|---|---|
| ISG15_iTRAQ1 | Scan_Id | 113 | 113, 117 | u_scan | WT-C-1 |
| ISG15_iTRAQ1 | Scan_Id | 114 | 113, 117 | u_scan | WT-AngII-1 |
| ISG15_iTRAQ1 | Scan_Id | 115 | 113, 117 | u_scan | ISG15-C-1 |
| ISG15_iTRAQ1 | Scan_Id | 116 | 113, 117 | u_scan | ISG15-AngII-1 |
| ISG15_iTRAQ1 | Scan_Id | 117 | 113, 117 | u_scan | WT-C-2 |
| ISG15_iTRAQ1 | Scan_Id | 118 | 113, 117 | u_scan | WT-AngII-2 |
| ISG15_iTRAQ1 | Scan_Id | 119 | 113, 117 | u_scan | ISG15-C-2 |
| ISG15_iTRAQ1 | Scan_Id | 121 | 113, 117 | u_scan | ISG15-AngII-2 |
| ISG15_iTRAQ2 | Scan_Id | 113 | 113, 117 | u_scan | WT-C-3 |
| ISG15_iTRAQ2 | Scan_Id | 114 | 113, 117 | u_scan | WT-AngII-3 |
| ISG15_iTRAQ2 | Scan_Id | 115 | 113, 117 | u_scan | ISG15-C-3 |
| ISG15_iTRAQ2 | Scan_Id | 116 | 113, 117 | u_scan | ISG15-AngII-3 |
| ISG15_iTRAQ2 | Scan_Id | 117 | 113, 117 | u_scan | WT-C-4 |
| ISG15_iTRAQ2 | Scan_Id | 118 | 113, 117 | u_scan | WT-AngII-4 |
| ISG15_iTRAQ2 | Scan_Id | 119 | 113, 117 | u_scan | ISG15-C-4 |
| ISG15_iTRAQ2 | Scan_Id | 121 | 113, 117 | u_scan | ISG15-AngII-4 |

*Figure S9. The LEVEL CREATOR task table for workflow 2.*

LEVEL CREATOR generates the *u_scan* (*uncalibrated scan*) files, which contain the scan identifiers (taken from the column “Scan_Id” in the “ID-q” table), the log2-ratios at the scan level Xs (as defined in the task table) and the *uncalibrated* weights Vs (which in the WSPP model are the intensities of the reporters in the Ratio numerator column) (*Figure S10*). The uncalibrated weights Vs are related to the quality of quantification (a higher weight implicates a more accurate quantification), but they are not associated to a statistical variance yet.

| Scan_Id | Xs_116_vs_113-117_Mean | Vs_116_vs_113-117_Mean |
|---|---|---|
| ISG15_iTRAQ1_v2-26378-2 | 0.166739599 | 354392.25 |
| ISG15_iTRAQ1_v2-61753-2 | 0.513614912 | 574721.75 |
| iTRAQ1_FR3-31890-3 | 0.055997478 | 51850.67578 |
| ISG15_iTRAQ1_v2-62222-2 | 0.311700218 | 939171.75 |
| iTRAQ1_FR2-28607-3 | 0.334866702 | 32351.34375 |
| iTRAQ1_FR1_20190705161217-23524-2 | 0.516515769 | 2680378.5 |

*Figure S10. Excerpt from one of the* u_scan *files generated by workflow 2 LEVEL CREATOR module showing element identifiers (left column), log2 ratios (center column), and statistical weights (right column).*

The LEVEL CALIBRATOR module calibrates the Vs weights by performing a *u_scan-to-peptide integration* and generates the *scan* (*calibrated scan*) files, which contain true, calibrated statistical weights (defined in the WSPP model as the inverse of the estimated individual scan variances) (*Figure S11***Error! Reference source not found.**, Top).

| Sample f | Lower level for | Higher level for ir | Name of calibrated level |
|---|---|---|---|
| * | u_scan | peptide | scan |

| Sample folder(s) | Lower level | Higher level | Output Sample folder | Tag | FDR |
|---|---|---|---|---|---|
| * | scan | peptide | | | |
| * | peptide | protein | | | |
| * | protein | proteinall | | | 0 |

*Figure S11. The LEVEL CALIBRATOR (Top) and INTEGRATE (Bottom) task tables for workflow 2.*

Note that LEVEL CALIBRATOR automatically generates a plot to supervise the accuracy of calibrations ("*_outGraph_VRank" png file) in each sample folder, which show whether the model is able to predict experimental scan variances as a function of the calibrated statistical weights (*Figure S12*).

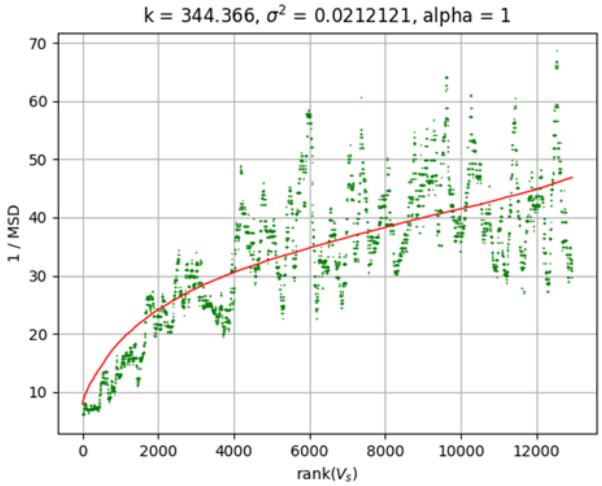

*Figure S12. Automatically generated graphs to supervise the accuracy of calibrations. These graphs represent 1/MSD versus the rank of Vs (scan weight, which in this level corresponds to reporter intensity). MSD is the Mean Squared Deviation of the scans vs the respective mean of the peptide they belong to. The scans are ordered by Vs and the MSD is calculated in a sliding window of 200 scans [2].*

As in the case of workflow 1, before performing the integrations, the relation tables have to be created with the RELS CREATOR module, which have a similar structure (*Figure S13*).

| Relation Table to be created | Column name of Lower level | Column name of Higher level | Column na | Table from which RT is extracted |
|---|---|---|---|---|
| u_scan2peptide | Scan_Id | Pep_Id | | |
| scan2peptide | Scan_Id | Pep_Id | | |
| peptide2protein | Pep_Id | Master Protein Accessions | | |
| protein2proteinall | Master Protein Accessions | [1] | | |
| protein2category | Protein | Category | | {PATH}/uniprot-MusMusculus_GOanotationDavid_enero2018_rels_v8_Protein2Category.txt |
| category2categoryall | Category | [1] | | {PATH}/uniprot-MusMusculus_GOanotationDavid_enero2018_rels_v8_Protein2Category.txt |

*Figure S13. The RELS CREATOR task table for workflow 2.*

The INTEGRATE module performs the *scan-to-peptide*, *peptide-to-protein* and *protein-to-proteinall* integrations according to the module task table (*Figure S11***Error! Reference source not found.**, Bottom). Note that for consistency all the files created for each sample are stored by default in the folder indicated in the task table of LEVEL CREATOR module, unless otherwise indicated in the Output Sample folder column.

Note that the INTEGRATE module automatically generates a plot to check the accuracy of the GIA integration model in each one of the integration steps. This is done by comparing the distribution of Z values with that of the null hypothesis (standard normal distribution) (see **Error! Reference source not found.** below, left panels). These graphs are stored in the corresponding sample folders (in "*_outGraph.png" files).

By default, iSanXoT removes integration outliers. To prevent the removal of outlier elements in the *protein-to-proteinall* integration, as these are just the proteins which are significantly altered, a *0 FDR* value was indicated in the INTEGRATE task table for this integration (Figure S11***Error! Reference source not found.***, Bottom).

Once protein levels are created, workflow 2 uses the NORCOMBINE basic module (*Figure S8*B) to integrate protein values from the four biological replicates in each group to produce integrated protein values per group that are stored in the folders WT-C, WT-AngII, ISG15-C and ISG15-AngII (*Figure S14*).

| Sample folders | Level | Norm | lowerNorm | Output Sample folder |
|---|---|---|---|---|
| WT-C-1 , WT-C-2 , WT-C-3 , WT-C-4 | protein | proteinall | lowerNormV | WT-C |
| WT-AngII-1 , WT-AngII-2 , WT-AngII-3 , WT-AngII-4 | protein | proteinall | lowerNormV | WT-AngII |
| ISG15-C-1 , ISG15-C-2 , ISG15-C-3 , ISG15-C-4 | protein | proteinall | lowerNormV | ISG15-C |
| ISG15-AngII-1 , ISG15-AngII-2 , ISG15-AngII-3 , ISG15-AngII-4 | protein | proteinall | lowerNormV | ISG15-AngII |

*Figure S14. The NORCOMBINE task table for workflow 2.*

The NORCOMBINE module integrates biological replicates within sample groups applying the GIA algorithm [1], which models the distribution of protein values around the average taking into account error propagation theory and estimates a global variance for the integration. The GIA algorithm assumes that the individual variances of all the lower elements (proteins) are affected by a global variance (which in this case arises from biological variability within the same group). While this assumption may not hold in all the cases, it can be easily checked by inspecting the test distributions. The NORCOMBINE module (like the INTEGRATE module) automatically generates graphs comparing the distribution of the integrated Z variables with those of the standard normal distribution. As shown in ***Error! Reference source not found.*** right, the distribution of protein Z values estimated by the model in the case of the ISG15-AngII group agree very well with the null hypothesis, demonstrating that the assumption of the model is a good approach to treat the biological variance of the samples within this group. Similar results were obtained in the other three groups (not shown).

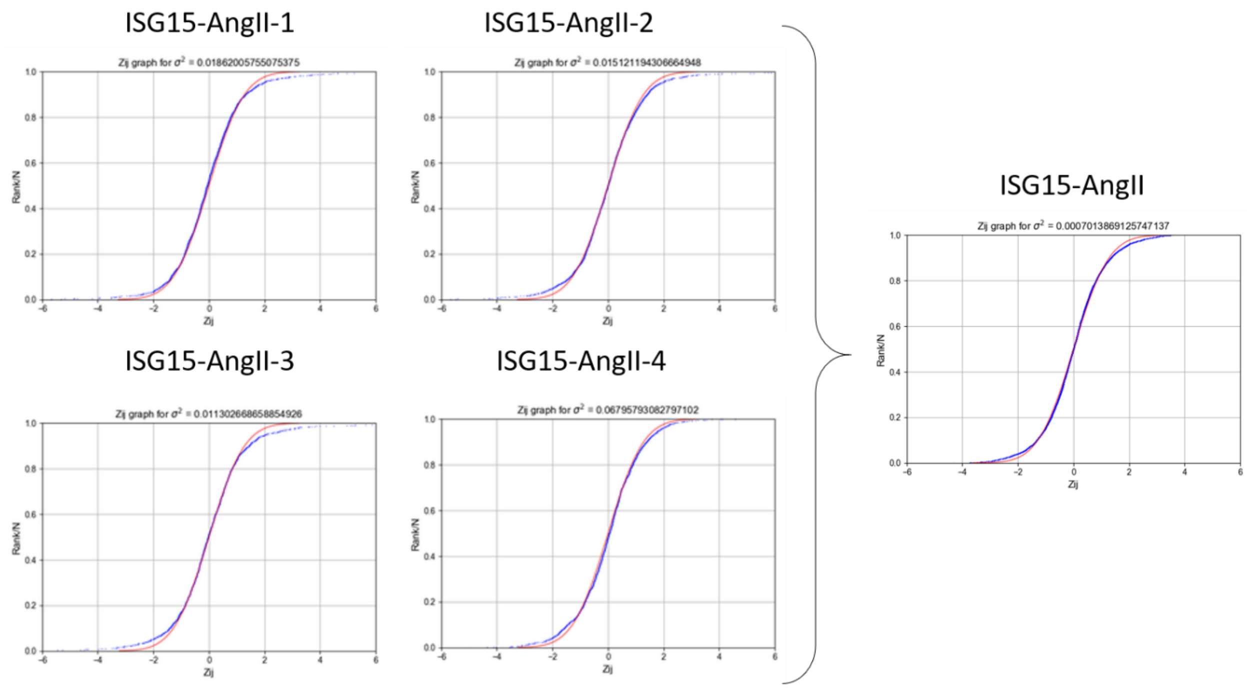

*Figure S15. Distribution of the standardized log2 protein ratios (Zqa) from the four individual ISG15-AngII VSMC samples (Left panel) showing how the WSPP model agrees well with the expected null distributions in the four cases, and from the integrated ISG15-AngII sample group obtained with the NORCOMBINE module (Right panel), showing how the GIA assumption of a global biological variance is a good approach to treat the biological variability of samples within this group. Red: null hypothesis (standard distribution); blue: experimental data.*

Note also that the NORCOMBINE module performs a weighted averaging from several samples and that from the good fitting to the null hypothesis it allows an accurate control over outliers (*Figure S15*). This unique approach allows the integration of protein values originating from unbalanced sample groups, distinct experiments or mass spectrometers, and even different labelling techniques (see for instance [2]).

The module task table indicates that samples were combined at the *protein* level using the *proteinall* level for normalization. This means that log2 protein ratios are firstly normalized by the grand mean before being integrated into an averaged protein value; this compensates for differences in protein load into each iTRAQ channel. Note that proteins could also be integrated to other levels (for instance organules, subcellular compartments, complexes, …), before being integrated by NORCOMBINE, allowing different kinds of normalizations. Finally, the column lowerNorm indicates which is the file that contains the normalized data, which usually are the *lowerNormV* files, previously generated by the INTEGRATE module. Further details can be found in the iSanXoT documentation.

The protein averages from the four biological sample groups are then used by the RATIOS basic module to calculate two ratios: *WT-AngIIvsWT-C*, where wild-type AngII-treated animals are compared to controls; and *ISG15-AngIIvsISG15-C*, where ISG15 AngII-treated animals are compared to ISG15 controls (*Figure S16*). The V method column allows the user to indicate the method used to assign a statistical weight to the log2ratios, which by default is the method called “max” (for further details see “RATIOS” module in the iSanXoT wiki, https://github.com/CNIC-Proteomics/iSanXoT/wiki).

| Ratio numerator c | Ratio denominator | Level | V Method | Output Sample folder |
|---|---|---|---|---|
| WT-AngII | WT-C | protein | max | WT-AngIIvsWT-C |
| ISG15-AngII | ISG15-C | protein | max | ISG15-AngIIvsISG15-C |

*Figure S16. The RATIOS task table for workflow 2.*

The last basic module executed in workflow 2, SBT (*Figure S17*), applies the SBT algorithm to the above defined comparisons for the detection of functional category changes originated by the coordinated behaviour of proteins. This is made as explained in workflow 1. The SBT module is more flexible since it allows to perform the triangle operations to any kind of level, not only proteins. In this case the triangle is made by the levels protein and category (*Figure S16*) and the corresponding grand mean.

| Sample folder(s) | Lower level | Intermediate level |
|---|---|---|
| WT-AngIIvsWT-C | protein | category |
| ISG15-AngIIvsISG15-C | protein | category |

*Figure S17. The SBT task table for workflow 2.*

Finally, the REPORT module (*Figure S18*) is used as in workflow 1 to generate tables with the protein and category data. In this case additional features of the REPORT module are used. The table “Npep2prot" is generated using an asterisk. This symbol is used by iSanXoT as a wildcard character to indicate that the results from all the samples containing the *peptide-to-protein* integration (i.e. ISG15-AngII-1, ISG15-AngII-2, ISG15-AngII-3, ISG15-AngII-4, ISG15-AngII, ISG15-C-1, ISG15-C-2, ISG15-C-3, ISG15-C-4, ISG15-C, and ISG15-AngIIvsISG15-C) are to be included in the table. However, the “Npep2prot_Quanprot_ISG15_filtered” and “Npep2prot_Quanprot_WT_filtered” tables include the protein changes (Zqa), the statistical significance (FDRqa) of these changes, and the number of peptides per protein only from the samples indicated in the *Sample folder(s)* column. The report for the ISG15 samples is filtered by Zqa to show the most extreme values (greater than 1 or less than -1) but only for the “ISG15-AngIIvsISG15-C” sample. Additional filters for the minimum number of peptides per protein are also used in these tables. The tables containing category values are filtered by Zca (greater than or equal to 2 or less than or equal to -2) and/or by the number of proteins per category (between 5 and 100).

| Sample folder(s) | Lower level | Higher level | Reported vars | Output report | Column headers to elimi | Merge with report | Ad | Filter |
|---|---|---|---|---|---|---|---|---|
| * | peptide | protein | n | Npep2prot | | | | |
| ISG15-AngII-1 , ISG15-AngII-2 , ISG15-AngII-3 , ISG15-AngII-4 , ISG15-AngII , ISG15-C-1 , ISG15-C-2 , ISG15-C-3 , ISG15-C-4 , ISG15-C , ISG15-AngIIvsISG15-C | protein | proteinall | Z , FDR | Npep2prot_Quanprot_ISG15_filtered | peptide | Npep2prot | | (Z_protein2proteinall@ISG15-AngIIvsISG15-C <= -1 \| Z_protein2proteinall@ISG15-AngIIvsISG15-C >= 1) & (n_peptide2protein >= 2) |
| WT-AngII-1 , WT-AngII-2 , WT-AngII-3 , WT-AngII-4 , WT-AngII , WT-C-1 , WT-C-2 , WT-C-3 , WT-C-4 , WT-C , WT-AngIIvsWT-C | protein | proteinall | Z , FDR | Npep2prot_Quanprot_WT_filtered | peptide | Npep2prot | | (n_peptide2protein > 4) |
| ISG15-AngIIvsISG15-C , WT-AngIIvsWT-C | protein | category | n | Nprot2cat | | | | |
| ISG15-AngIIvsISG15-C , WT-AngIIvsWT-C | category | categoryall | Z , FDR | Nprot2cat_Quancat_filtered | protein | Nprot2cat | | (Z_category2categoryall <= -2 \| Z_category2categoryall >= 2) & (n_protein2category >= 5) & (n_protein2category <= 100) |
| ISG15-AngIIvsISG15-C , WT-AngIIvsWT-C | protein | proteinall | Z , FDR | Nprot2cat_Quanprot | | Nprot2cat | | |
| ISG15-AngIIvsISG15-C , WT-AngIIvsWT-C | category | categoryall | Z , FDR | Nprot2cat_Quancat_Quanprot_filtered | | Nprot2cat_Quanprot | | (n_protein2category >= 5) & (n_protein2category <= 100) |

*Figure S18. The REPORT task table for workflow 2.*

The tables generated by REPORT can be used to generate heatmaps showing the most relevant protein abundance changes (*Figure S19*). As previously shown [3], iSanXoT analysis revealed a coordinated alteration of proteins implicated in cardiovascular function, extracellular matrix and remodeling, and vascular redox state in aortic tissue from AngII-infused ISG15-KO mice (*Figure S20A*). The coordinated protein behavior from some of the altered categories can be analyzed in the sigmoid plots (*Figure S20B*).

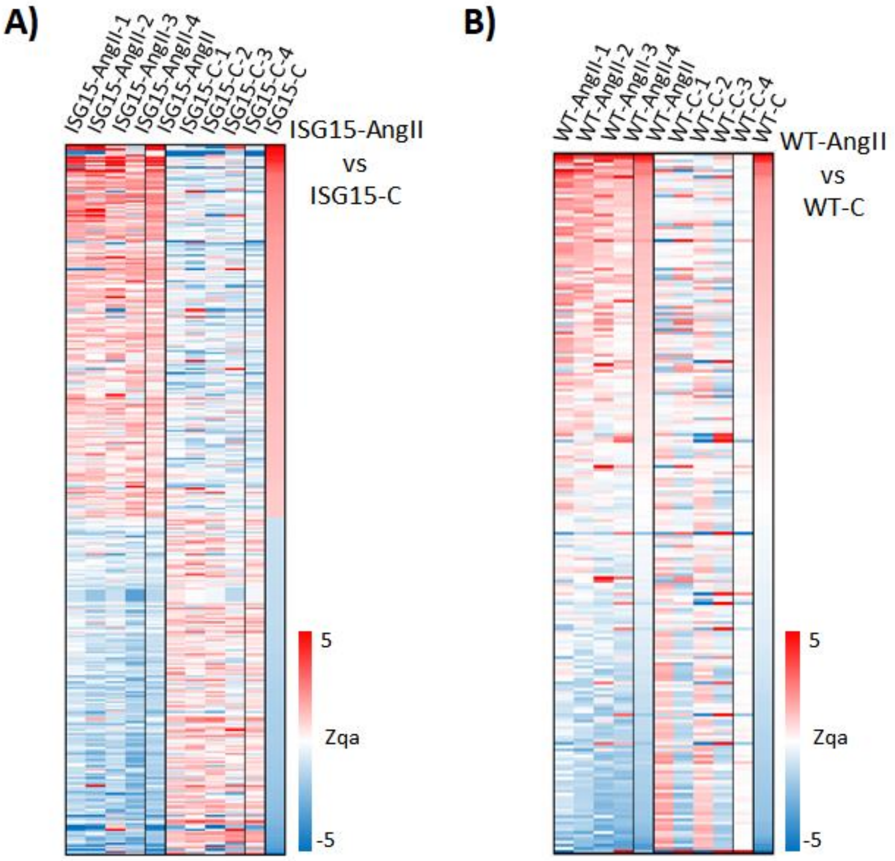

*Figure S19. Differential abundance of functional proteins revealed by workflow 2. The heatmap (A) for proteins (Zqa) is based on the "Npep2prot_Quanprot_ISG15_filtered" REPORT table. The heatmap (B)*

*displays the proteins (Zqa) for the WT samples using the "Npep2prot_Quanprot_WT_filtered" REPORT table.*

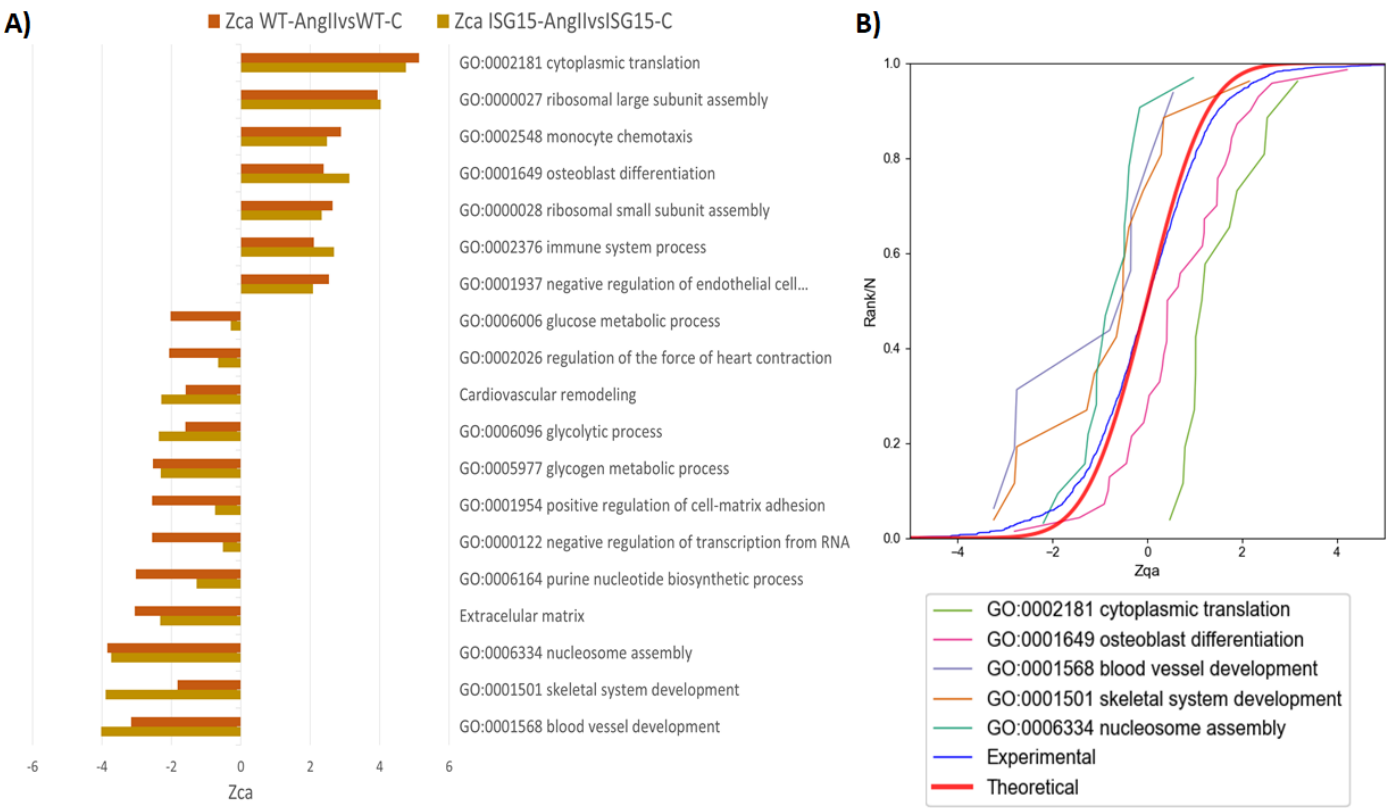

*Figure S20. Functional category changes arising from coordinated protein behavior. A) Bar graph for functional categories (Zca) constructed from the "Nprot2cat_Quancat_filtered" REPORT table. B) The distributions of the standardized log2 protein ratios (Zqa) are shown for some of the functional categories that are significantly down-regulated (Left) or up-regulated (Right). The data to create the sigmoid curves are taken from the "Nprot2cat_Quancat_Quanprot_filtered" REPORT table.*

### Workflow execution

The workflow template and input files that are needed to execute this workflow can be downloaded from https://github.com/CNIC-Proteomics/iSanXoT/wiki/studies/cases/templates/WSPP_NORCOM_RATIOS_SBT.zip. See the *Importing a workflow template* Section below for detailed instructions.

### Workflow 3: Quantification of posttranslationally modified peptides in a labeled experiment

#### Experimental

This workflow was used to quantitate reversibly oxidized Cys peptides in mouse embryonic fibroblast (MEF) preparations subjected to chemical oxidation with diamide, an experiment that served to illustrate the comparative performance of on-filter (FASILOX) and in-gel (GELSILOX) approaches to study the thiol redox proteome [4]. These techniques introduced a differential label on Cys residues, depending on their oxidation state, producing two separate populations of reduced and oxidized Cys-containing peptides. MEF samples were incubated with diamide (treated group), or PBS (control group) and the resulting peptides were isobaricaly labeled with iTRAQ 8-plex (four biological replicates per condition). The workflow detects statistically significant abundance changes in peptides containing modified Cys residues.

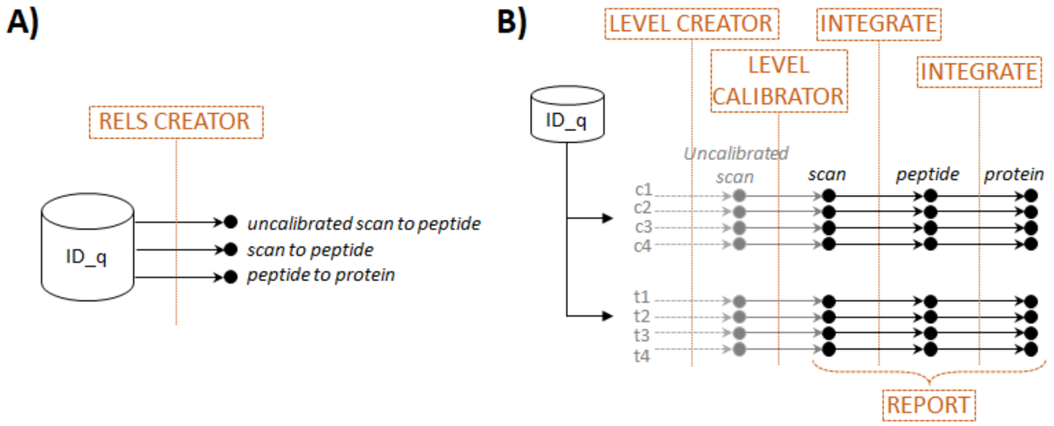

*Figure S21. Scheme of workflow 3 (quantification of posttranslationally modified peptides in a labeled experiment) showing module components: RELS CREATOR (A) and LEVEL CREATOR, LEVEL CALIBRATOR, INTEGRATE, and REPORT (B).*

#### Workflow operation

Workflow 3 comprises the basic modules LEVEL CREATOR, LEVEL CALIBRATOR, and INTEGRATE, as well as the RELS CREATOR and REPORT modules (*Figure S21*) and is very similar to workflow 2. LEVEL CREATOR was used to design the ratios and to generate the level files, sample folders and log2 ratios indicated in the corresponding task table (*Figure S22* and *Figure S23*). LEVEL CALIBRATOR was used to calibrate statistical weights (*Figure S24*, top) and INTEGRATOR to integrate from scan to peptide and from peptide to protein (*Figure S24*, bottom).

| Experiment | Identifier column header | Ratio numerator column | Ratio denominator column(s) | Level to be created | Output Sample folder |
|---|---|---|---|---|---|
| JAL_FASILOX_iTR_ALL | ScanIdentifier | q_reporterIon_113 | q_reporterIon_117 , q_reporterIon_118 , q_reporterIon_119 , q_reporterIon_121 | uscan | c1 |
| JAL_FASILOX_iTR_ALL | ScanIdentifier | q_reporterIon_114 | q_reporterIon_117 , q_reporterIon_118 , q_reporterIon_119 , q_reporterIon_121 | uscan | c2 |
| JAL_FASILOX_iTR_ALL | ScanIdentifier | q_reporterIon_115 | q_reporterIon_117 , q_reporterIon_118 , q_reporterIon_119 , q_reporterIon_121 | uscan | c3 |
| JAL_FASILOX_iTR_ALL | ScanIdentifier | q_reporterIon_116 | q_reporterIon_117 , q_reporterIon_118 , q_reporterIon_119 , q_reporterIon_121 | uscan | c4 |
| JAL_FASILOX_iTR_ALL | ScanIdentifier | q_reporterIon_117 | q_reporterIon_113 , q_reporterIon_114 , q_reporterIon_115 , q_reporterIon_116 | uscan | t1 |
| JAL_FASILOX_iTR_ALL | ScanIdentifier | q_reporterIon_118 | q_reporterIon_113 , q_reporterIon_114 , q_reporterIon_115 , q_reporterIon_116 | uscan | t2 |
| JAL_FASILOX_iTR_ALL | ScanIdentifier | q_reporterIon_119 | q_reporterIon_113 , q_reporterIon_114 , q_reporterIon_115 , q_reporterIon_116 | uscan | t3 |
| JAL_FASILOX_iTR_ALL | ScanIdentifier | q_reporterIon_121 | q_reporterIon_113 , q_reporterIon_114 , q_reporterIon_115 , q_reporterIon_116 | uscan | t4 |

*Figure S22. The LEVEL CREATOR task table for workflow 3.*

| ScanIdentifier | Xs_q_reporterIon_113_vs_q_reporterIon_1 | Vs_q_reporterIon_113_vs_q_reporterIon_1 |
|---|---|---|
| JAL_FASILOX_iTR_ALL.raw-14205-3 | -0.760406365 | 2154536.848 |
| JAL_FASILOX_iTR_ALL.raw-19883-2 | -0.64797195 | 475243.9143 |
| JAL_FASILOX_iTR_ALL.raw-51554-3 | -0.567309329 | 630711.4777 |
| JAL_FASILOX_iTR_ALL.raw-77608-4 | -0.620612557 | 826786.5206 |
| JAL_FASILOX_iTR_ALL.raw-13670-2 | -0.42826962 | 445258.3775 |
| JAL_FASILOX_iTR_ALL.raw-50717-2 | -0.490418129 | 324232.4633 |

*Figure S23. Excerpt from one of the* uscan *files generated by workflow 3 LEVEL CREATOR module showing element identifiers (left column), log2 ratios (center column) and statistical weights (right column).*

The only difference with workflow 2 lies in the INTEGRATE command used for the integration *peptide-to-protein*. INTEGRATE can use a modified version of the GIA algorithm for the quantitative analysis of post-translational modifications (PTM) that includes a third column containing *tags* in the relation tables, as described [5]. In this workflow the advanced option of INTEGRATE was activated to display the *Tag* column, which is used to include only the peptides which are tagged in the relation table with the text *"Not modified"* when calculating the protein averages (*Figure S24*). An example of tagged *peptide2protein* relation table is shown in *Figure S25.* Proteins are thus quantified using only peptides which are not modified in Cys. However, although these Cys peptides do not contribute to protein averages, they are assigned a *Zpq* value, which serves to evaluate whether they deviate significantly from the expected distribution of peptides around their protein averages [4]. If the deviation is statistically significant it can be concluded that there is a change in abundance of the posttranslational modification in relation to the protein it comes from. This philosophy can be extended to any other kind of PTM.

| Sample folder(s) | Lower level for integration | Higher level for integration | Name of calibrated level |
|---|---|---|---|
| * | uscan | peptide | scan |

| Sample folder(s) | Lower level | Higher level | Output Sample folder | Tag |
|---|---|---|---|---|
| * | scan | peptide | | |
| * | peptide | protein | | Not modified |

*Figure S24. The LEVEL CALIBRATOR (Top) and INTEGRATE (Bottom) task tables for workflow 3.*

| protein | peptide | Modifications |
|---|---|---|
| >tr\|E9Q7Q3\|E9Q7Q3_MOUSE Tropom | EQAEAEVASLNRR | Not modified |
| >tr\|E9Q7Q3\|E9Q7Q3_MOUSE Tropom | EQAEAEVASLNR | Not modified |
| >sp\|Q8BTM8\|FLNA_MOUSE Filamin- | SNFTVDC@SK{ | Reduced-Cys peptides |
| >sp\|P48962\|ADT1_MOUSE ADP/ATP | DFLAGGIAAAVSK{ | Not modified |
| >sp\|Q8VDN2\|AT1A1_MOUSE Sodium/ | NLEAVETLGSTSTIC@SDK{ | Reduced-Cys peptides |
| >sp\|P20152\|VIME_MOUSE Vimentin | QVQSLTC#EVDALK{ | Oxidized-Cys peptides |
| >sp\|Q9CPY7\|AMPL_MOUSE Cytosol | QVIDC@QLADVNNLGK{ | Reduced-Cys peptides |
| >sp\|Q501J6\|DDX17_MOUSE Probabl | GVEIC@IATPGR | Reduced-Cys peptides |
| >sp\|Q9CZ44\|NSF1C_MOUSE NSFL1 c | LGSTAPQVLNTSSPAQQAENEAK{ | Not modified |
| >sp\|B2RSH2\|GNAI1_MOUSE Guanine | TTGIVETHFTFK{ | Not modified |

*Figure S25. Excerpt from the peptide2protein relation table used to integrate peptides to proteins. Note the presence of a third column used to tag Cys-containing peptides, which will be excluded from the calculation of protein averages in the peptide-to-protein integration.*

iSanXoT allows to automatically generate relation tables containing tags, which are taken from the "ID-q" table. For this end, RELS CREATOR uses a specific option (*Figure S26*). In this particular case, this option makes RELS CREATOR to look into the "ID-q" table for the column with the header *Modifications* and translate its content into the third column of the *peptide2protein* relation table. In the "ID-q" table used in this case, the peptide containing modified Cys residues were labeled as "*Reduced-Cys peptides*" and "*Oxidized-Cys peptides*" depending on the type of modification. These tags are located in the relation table by RELS CREATOR (*Figure S25* and *Figure S26*). iSanXoT allows to use any tag created by searching engines or defined by the user, with the only condition that the tag indicated in the INTEGRATE command must match the tag in the third column of the relation table (*Figure S24*, bottom).

| Relation Table to be created | Column name of Lower level | Column name of Higher level | Column name of 3rd column |
|---|---|---|---|
| uscan2peptide | ScanIdentifier | Sequence | |
| scan2peptide | ScanIdentifier | Sequence | |
| peptide2protein | Sequence | FASTAshort | Modifications |

*Figure S26. The RELS CREATOR task table for workflow 3.*

Finally, the REPORT module collects the statistical variables generated by the *peptide-to-protein* integration for all the samples (*c1, c2, c3, c4, t1, t2, t3,* and *t4*), as prompted by the asterisk (*Figure S27A*). The REPORT commands are similar to those used to generate the protein tables in workflow 1, with the difference that peptide values are tabulated instead of protein values, together with the number of scans per peptide, instead of the number of peptides per protein. The FDR at the peptide level allows to detect statistically significant changes in PTM. This REPORT also generates a second filtered peptide table containing the peptides with reduced Cys with the most extreme abundance changes. This table was used to generate a heatmap (*Figure S27B*).

**A)**

| Sample fo | Lower level | Higher level | Reported vars | Output report | Column headers to elim | Merge with report | Ad | Filter |
|---|---|---|---|---|---|---|---|---|
| * | scan | peptide | n | Nscan2pep | | | | |
| * | peptide | protein | Z , FDR, tags | Nscan2pep_Quanpepprot | scan | Nscan2pep | | |
| * | peptide | protein | Z , tags | Nscan2pep_Quanpepprot_filtered | scan | Nscan2pep | | (tags_peptide2protein == "Reduced-Cys peptides") & (Z_peptide2protein >= 2.5 \| Z_peptide2protein <= -2.5) |

**B)**

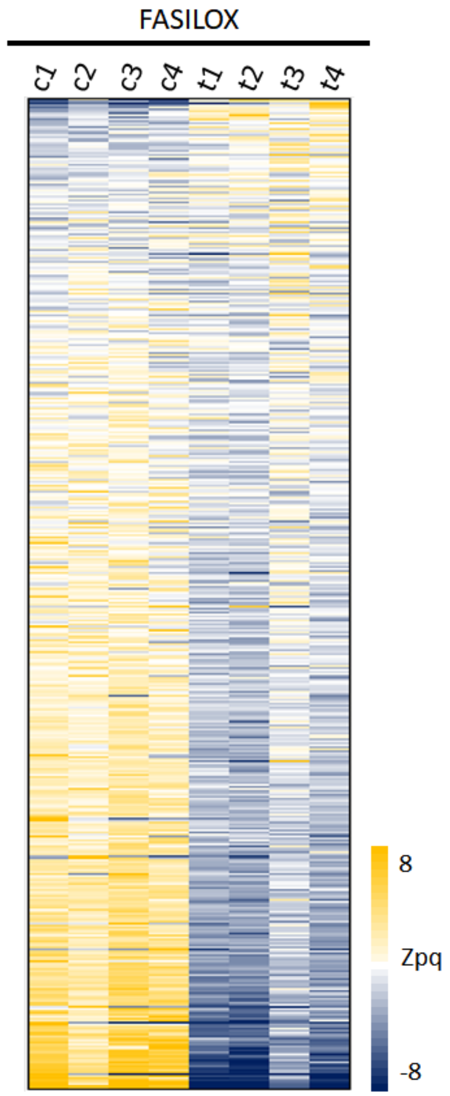

*Figure S27. (A) The task table for Workflow 3 is in the REPORT module. (B) Relative abundance of Cys-containing peptides in MEF samples measured by peptide log2 ratios expressed in units of standard deviation corrected by the protein mean (Zpq). The data for the heatmap was generated from the "Nscan2pep_Quanpepprot_filtered" report table.*

Of note, the peptides included in the integration followed a standard distribution in the eight samples, as shown in blue on *Figure S28* for *t1*, *t2*, *c1*, and *c2* samples. This evidences that the error distribution at the peptide level could be accurately modeled using the GIA algorithm. In addition, the treatment produced a generalized increase in the abundance of oxidized Cys-containing peptides (orange curves), with concomitant decrease in the abundance of reduced Cys-containing peptides (green curves). Consistently, the opposite changes were observed in the controls.

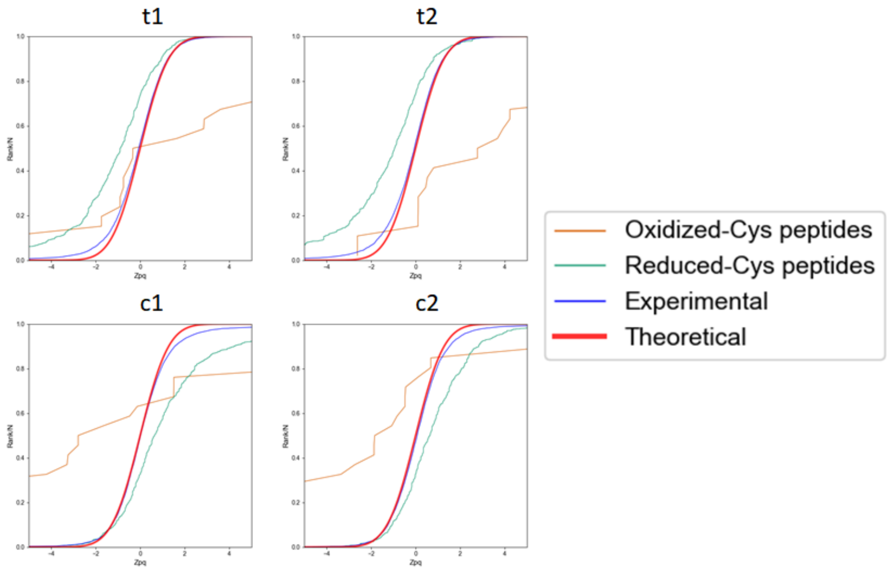

*Figure S28. Distribution of the standardized variable at the peptide level (Zpq) in control MEF samples (t1, t2, c1 and c2) for all the peptides quantitated (blue) and the oxidized (orange) and reduced (green) Cys-containing peptide subpopulations. The theoretical normal distribution N(0,1) is shown in red. Positive/negative Zpq values indicate increased/decreased peptide abundance with respect to the average. These sigmoidal curves were created from the "Nscan2pep_Quanpepprot" table generated by REPORT.*

#### Workflow execution

The workflow template and input files that are needed to execute this workflow can be downloaded from

https://github.com/CNIC-Proteomics/iSanXoT/wiki/studies/cases/templates/WSPP_PTM.zip

See the *Importing a workflow template* Section below for detailed instructions.

### Workflow 4: Label-free quantification

#### Experimental

This workflow was used to analyze quantitative data obtained in a multicenter study aimed at evaluating different bioinformatics tools [6]. The authors generated two hybrid proteome test samples consisting of tryptic digests of human, yeast and *Escherichia coli* proteins mixed in two different defined proportions (13:3:4 for *A* samples and 13:6:1 for *B* samples). Quadruplicate peptide samples were analyzed by LC-MS/MS, after which peptide identification and quantification were carried out with several software packages. In this example we show how to process the data obtained by MaxQuant [7] (see Section below to learn how data from MaxQuant and other softwares can be adapted to be used with iSanXoT).

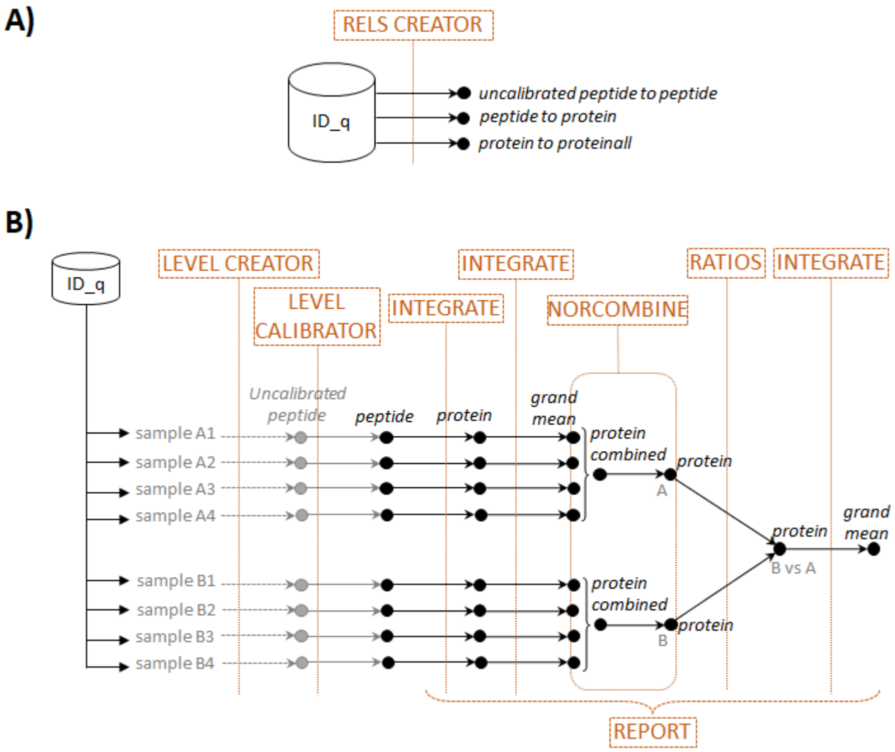

*Figure S29. Scheme of workflow 4 (label-free quantification) showing module components: RELS CREATOR (A) and LEVEL CREATOR, LEVEL CALIBRATOR, INTEGRATE, NORCOMBINE, and REPORT (B).*

#### Workflow operation

Workflow 4 includes the basic iSanXoT modules LEVEL CREATOR, LEVEL CALIBRATOR, INTEGRATE, NORCOMBINE, and RATIOS along with the REPORT and RELS CREATOR modules (*Figure S29*). The starting module, LEVEL CREATOR, generates the level files, sample folders and log2 ratios indicated in the corresponding task (*Figure S30*) based on the quantitative data at the peptide level obtained with MaxQuant for replicate *A*- and *B*-type samples. In this example we used as denominator of the log2ratio the average of peptide intensities across all the samples. In this case, however, the averages of the four

*A*-type and the four *B*-type samples are first calculated separately (as indicated by the square brackets), then the average of the two averaged values is calculated (as indicated by the comma). This ensures that no log2-ratio is calculated when the four values are missing in either the *A* or the *B* sample group. This module generates uncalibrated files at the peptide level (*u_peptide*) (*Figure S31*).

| Experiment | Identifier column header | Ratio numerator column | Ratio denominator column(s) | Level to be created | Output Sample folder |
|---|---|---|---|---|---|
| | Peptide_Id | Intensity B_01 | [Intensity A_01 , Intensity A_02 , Intensity A_03 , Intensity A_04] , [Intensity B_01 , Intensity B_02 , Intensity B_03 , Intensity B_04] | u_peptide | b1 |
| | Peptide_Id | Intensity B_02 | [Intensity A_01 , Intensity A_02 , Intensity A_03 , Intensity A_04] , [Intensity B_01 , Intensity B_02 , Intensity B_03 , Intensity B_04] | u_peptide | b2 |
| | Peptide_Id | Intensity B_03 | [Intensity A_01 , Intensity A_02 , Intensity A_03 , Intensity A_04] , [Intensity B_01 , Intensity B_02 , Intensity B_03 , Intensity B_04] | u_peptide | b3 |
| | Peptide_Id | Intensity B_04 | [Intensity A_01 , Intensity A_02 , Intensity A_03 , Intensity A_04] , [Intensity B_01 , Intensity B_02 , Intensity B_03 , Intensity B_04] | u_peptide | b4 |
| | Peptide_Id | Intensity A_01 | [Intensity A_01 , Intensity A_02 , Intensity A_03 , Intensity A_04] , [Intensity B_01 , Intensity B_02 , Intensity B_03 , Intensity B_04] | u_peptide | a1 |
| | Peptide_Id | Intensity A_02 | [Intensity A_01 , Intensity A_02 , Intensity A_03 , Intensity A_04] , [Intensity B_01 , Intensity B_02 , Intensity B_03 , Intensity B_04] | u_peptide | a2 |
| | Peptide_Id | Intensity A_03 | [Intensity A_01 , Intensity A_02 , Intensity A_03 , Intensity A_04] , [Intensity B_01 , Intensity B_02 , Intensity B_03 , Intensity B_04] | u_peptide | a3 |
| | Peptide_Id | Intensity A_04 | [Intensity A_01 , Intensity A_02 , Intensity A_03 , Intensity A_04] , [Intensity B_01 , Intensity B_02 , Intensity B_03 , Intensity B_04] | u_peptide | a4 |

*Figure S30. The LEVEL CREATOR task table for workflow 4.*

| Peptide_Id | Xs_Intensity A_01_vs_Mean_Intensity | Vs_Intensity A_01_vs_Mean_Intensity |
|---|---|---|
| 6988_EALQSDWLPFELLASGGQK | 0.072239228 | 2378200 |
| 6990_EALTYDGALLGDR | 0.062201152 | 144200000 |
| 6991_EALVDTLTGILSPVQEVR | 0.354203027 | 43264000 |
| 6993_EAMECSDVIWQR | -0.435525774 | 8966841.667 |
| 6998_EAMGIYISTLK | 0.106526482 | 90203000 |
| 7000_EAMNDPLLER | 0.015057358 | 43534000 |

*Figure S31. Excerpt from one of the u_peptide files generated by workflow 4 LEVEL CREATOR module showing element identifiers (left column), log2 ratios (center column) and uncalibrated statistical weights (right column).*

The *u_peptide* level files are then calibrated with the LEVEL CALIBRATOR module by performing an integration to the protein level (*Figure S32, Top*), generating calibrated *peptide* level files. The *peptide-to-protein* and *protein-to-proteinall* integrations are then performed by the INTEGRATE module according to the module task table (*Figure S32*, Bottom).

Note that in this example the advanced option of INTEGRATE was activated to use the Tag column, so that only the proteins containing the *Homo sapiens* tag are used in the protein-to-proteinall integrations (*Figure S32, Bottom*). Restricting the integration to human proteins serves for two purposes: a) the normalization is done by the grand mean of the human proteins and is not affected by the presence of yeast or *E. coli* proteins. b) only human proteins are used to estimate the variance of the *protein-to-protein* all integration, avoiding the effect of yeast and E. coli proteins, whose ratios have a large deviation from the mean. Note that this procedure does not eliminate yeast or *E. coli* proteins from the normalized files that are later used by the NORCOMBINE module (see below).

| Sample folder(s) | Lower level for integration | Higher level for integration | Name of calibrated level |
|---|---|---|---|
| * | u_peptide | protein | peptide |

| Sample folder(s) | Lower level | Higher level | Output Sample folder | Tag | FDR |
|---|---|---|---|---|---|
| a1 , a2 , a3 , a4 , b1 , b2 , b3 , b4 | peptide | protein | | | |
| a1 , a2 , a3 , a4 , b1 , b2 , b3 , b4 | protein | proteinall | | Homo sapiens | 0 |
| B_vs_A | protein | proteinall | | Homo sapiens | 0 |

*Figure S32. The LEVEL CALIBRATOR (Top) and INTEGRATE (Bottom) task tables for workflow 4.*

As in workflow 3, the *protein-to-proteinall* relation table must contain a third column tagging the species from which each protein comes from (*Figure S33*). Note that the tag indicating the human proteins matches the tag indicated in INTEGRATE (*Figure S32, Bottom*).

| proteinall | protein | Species |
|---|---|---|
| 1 | B7UM99 | Escherichia coli |
| 1 | P0ACF8 | Escherichia coli |
| 1 | P24232 | Escherichia coli |
| 1 | P18440 | Homo sapiens |
| 1 | P01920 | Homo sapiens |
| 1 | O75147 | Homo sapiens |
| 1 | P33302 | Saccharomyces cerevisiae |
| 1 | P22147 | Saccharomyces cerevisiae |
| 1 | P06169 | Saccharomyces cerevisiae |

*Figure S33. Excerpt from the protein2proteinall workflow 4 relation table that links proteins to a constant value representing the protein grand mean. Note the use of a third column to tag proteins with their corresponding species for later species-specific protein-to-proteinall integration.*

The *protein-to-proteinall* relation table is automatically created by the RELS CREATOR module, which takes this information from a file (HumanSaccEcoliPME12_divide_by_species.tsv) created by the user. The file contains the relationship between protein identifiers (*Protein_Id* column header) and the species (*Species* column header) they come from.

| Relation Table to be created | Column name of Lower level | Column name of Higher level | Column name of 3rd column | Table from which RT is extracted |
|---|---|---|---|---|
| u_peptide2protein | Peptide_Id | Proteins | | |
| peptide2protein | Peptide_Id | Proteins | | |
| protein2proteinall | Protein_Id | [1] | Species | {PATH}/HumanSaccEcoliPME12_divide_by_species.tsv |

*Figure S34. The RELS CREATOR task table for workflow 4.*

Next, the normalized data at the *protein* level from the four replicates from each sample are combined into samples *A* and *B*, respectively, using the NORCOMBINE basic module (*Figure S35*, Top). To compare these two samples, new log2 ratios and statistical weights are calculated using the RATIOS basic module (*Figure S35*, Bottom). Finally, a *protein-to-proteinall* integration is carried out for the newly-generated *B_vs_A* sample by the module INTEGRATE (*Figure S32*, Bottom), using again the *Homo sapiens* tag.

| Sample folders | Level | Norm | lowerNorm | Output Sample folder |
|---|---|---|---|---|
| b1 , b2 , b3 , b4 | protein | proteinall | lowerNormV | B |
| a1 , a2 , a3 , a4 | protein | proteinall | lowerNormV | A |

| Numerator Sample folder | Denominator Sample folder(s) | Level | V Method | Output Sample folder |
|---|---|---|---|---|
| B | A | protein | avg | B_vs_A |

*Figure S35. The NORCOMBINE (Top) and RATIOS (Bottom) task tables for workflow 4.*

The REPORT module used in this workflow tabulates the data at protein level together with the number of peptides per protein, as in previous workflows (*Figure S36*). In this example, besides *Z* and *FDR*, the table includes the log2ratios of proteins from all the samples (*Xinf* from the *protein-to-proteinall* integration, to which we also refer as *Xq*), the grand mean (*Xsup* from the *protein-to-proteinall integration,* or *Xa*), the statistical weights (*Vinf* or *Vq*) and the species each protein belongs to (*tags*). The grand mean, which is used to normalize log2-ratios, and the statistical weights can be used to construct plots such as in *Figure S37* (see below).

| Sample folder(s) | Lower level | Higher level | Reported vars | Output report | Column headers to eliminate | Merge with report |
|---|---|---|---|---|---|---|
| * | peptide | protein | n | Npep2prot | | |
| * | protein | proteinall | Xinf, Xsup, Vinf , Z , FDR , tags | Npep2prot_Quanprot | peptide | Npep2prot |

*Figure S36. The REPORT module task table for workflow 4.*

It is noteworthy that variance modelling, normalization, standardization and statistical weighting, according to the GIA algorithm, are performed automatically, without data filtering, pre-processing or missing value imputation [6]***Error! Reference source not found.***, even in a situation where numerous proteins have highly imbalanced data. Moreover, the sigmoid plots automatically generated in each one of the integrations performed by INTEGRATE clearly demonstrate that the GIA algorithm accurately predicts the distribution of peptide quantifications around their proteins (*Figure S37A*) and of protein quantifications around the grand mean (*Figure S37B*). These results demonstrate that this statistical model is very suitable for the analysis of label-free data.

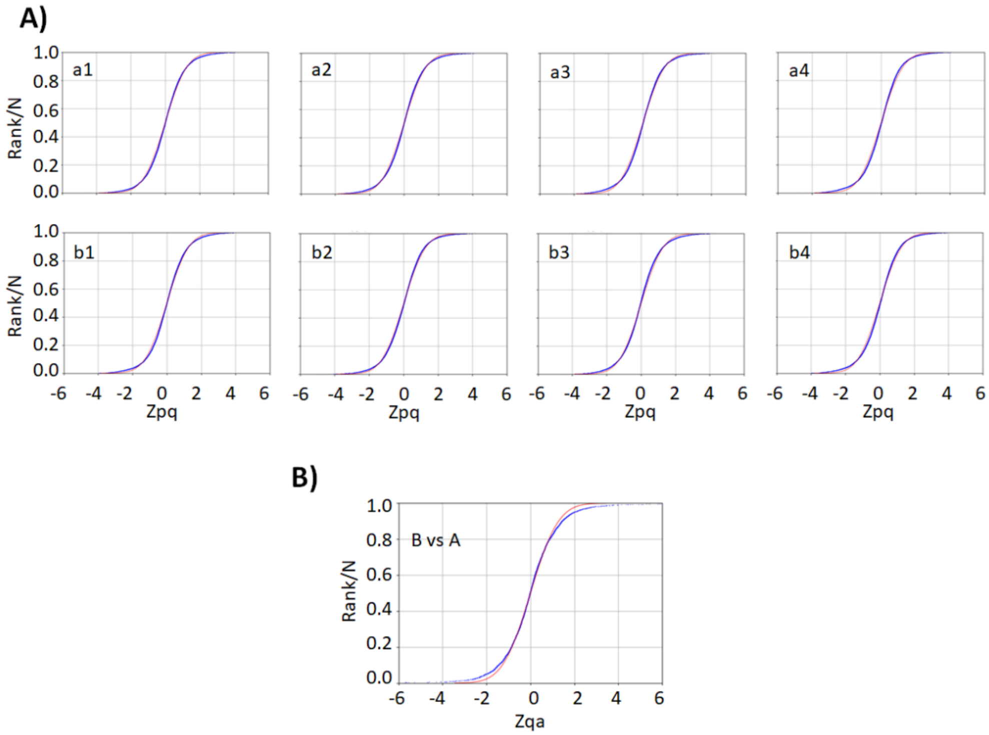

*Figure S37. Distribution of the standardized variable at the peptide (Zpq) and protein (Zqa) levels for label-free data analyzed with iSanXoT. A) Zpq distribution for the eight individual A-type and B-type samples. B) Zqa distribution for the B-type vs A-type comparison. Red: null hypothesis (standard distribution); blue: experimental data.*

The combined statistics *B_vs_A* also shows how human, yeast and bacterial proteins distribute around the expected 0, 1, and -2 log2- values (corresponding to 1-, 2- and 0.25-fold changes) (*Figure S38)*, and how protein quantifications with higher statistical weights are more accurate. This plot also confirms how iSanXoT provides highly accurate quantitative results in a fully automated fashion.

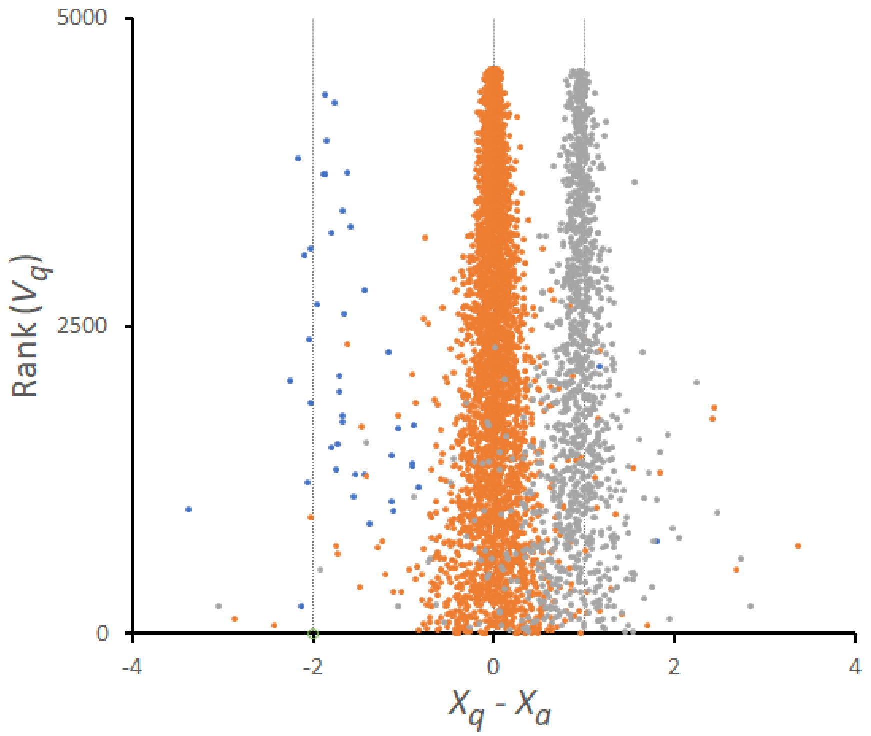

*Figure S38. Quantification of human (orange), yeast (grey) and bacterial (blue) proteins according to the combined statistics B_vs_A. Shown are log2-ratios normalized by the grand mean (Xq – Xa or Xinf - Xsup). This plot was generated from the table "Npep2prot_Quanprot".*

#### Workflow execution

The workflow template and input files that are needed to execute this workflow can be downloaded from

https://github.com/CNIC-Proteomics/iSanXoT/wiki/studies/cases/templates/WPP_LabelFree.zip

See the *Importing a workflow template* Section below for detailed instructions.

### Importing a workflow template

In this section we will provide instructions to execute the workflow examples and to import workflows that were previously created with iSanXoT to be reused in other projects. We will use as example the first workflow described in the previous section.

Start by downloading workflow 1 template and input files from the iSanXoT wiki (https://github.com/CNIC-Proteomics/iSanXoT/wiki/studies/workflows/WSPP-SBT.zip). Then extract the files included in the compressed archive to create a folder named WSPP-SBT. Check that the WSPP-SBT folder has been created in your file system. Then proceed as follows:

- Open the iSanXoT application by double-clicking the application icon (*Figure S39*).

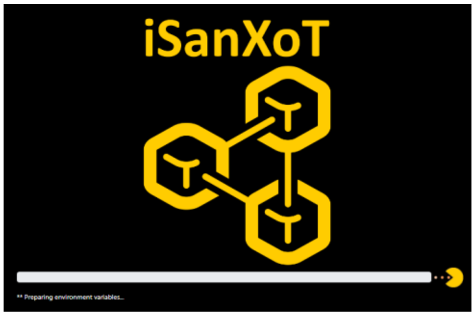

*Figure S39. The iSanXoT startup message.*

- Choose *New Project* from the *Project* menu (*Figure S40*).

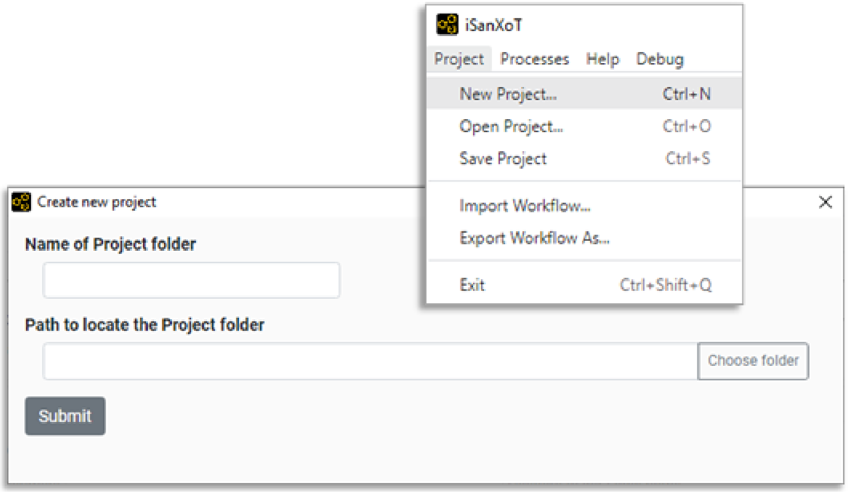

*Figure S40. Create New Project.*

- Provide a name of your choice for the project folder and indicate a path to locate this folder, then click the *Submit* button (*Figure S40*).
- Choose *Import Workflow* from the *Project* menu (*Figure S41)* and select the folder WSPP-SBT

created before (or any other iSanXoT project folder from which you want to import the workflow).

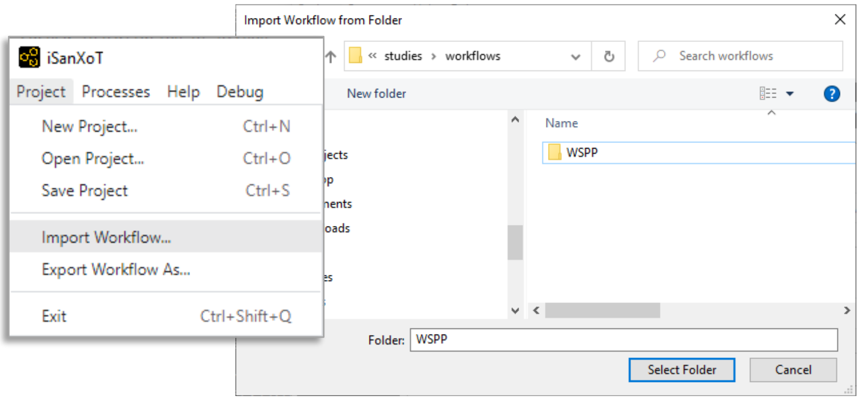

*Figure S41. Importing a preexisting iSanXoT workflow to the newly-created project.*

- Inspect the WSPP-SBT task table (in the Compound modules tab), the RELS CREATOR task table (in the Relation tables tab) and the REPORT task table (in the Reports tab) to check that the tables indicated in Fig. S2, S3 and S5 have been correctly loaded. Note that if a different template is imported, only the corresponding task tables will be loaded.

- Now click on *Choose identification file* and select “ID-q.tsv” in the WSPP-SBT folder (*Figure S42*). Or, alternatively, select the desired identification/quantification table with which this workflow is to be executed. Section 3 below shows how to prepare the “ID-q” file based on the output from a variety of proteomics pipelines. Bear in mind that the tasks defined in the LEVEL CREATOR and RELS CREATOR modules have to match the samples and column names from the specific “ID-q” file used.

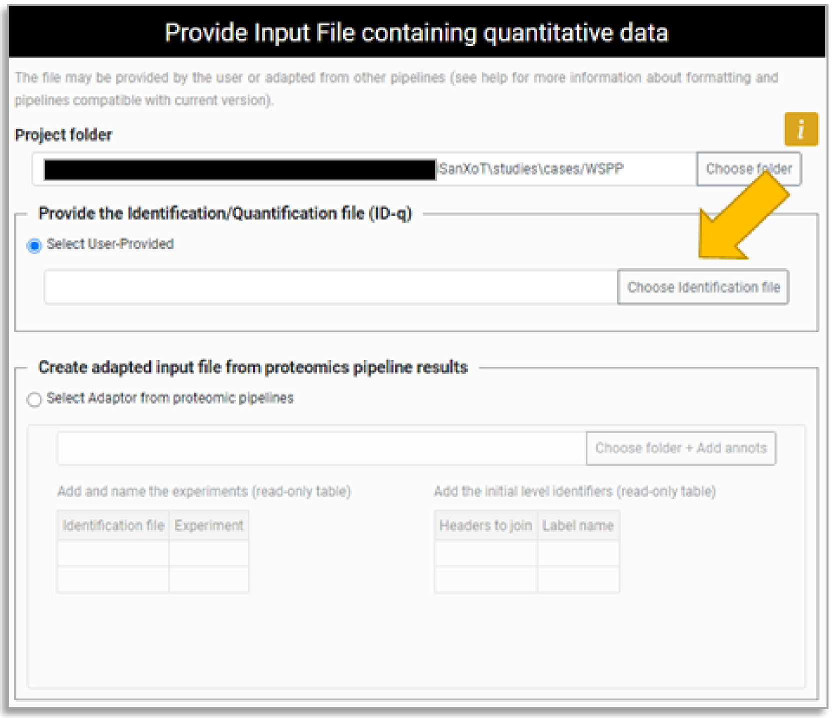

*Figure S42. Choosing the identification/quantification (ID-q) file for the newly-created project.*

- Select *Save Project* from the *Project* menu to save the changes or directly press the *Save and Run* button to save and execute the current workflow.

### Creating the identification/quantification file from proteomics pipelines

iSanXoT requires an identification/quantification file in tsv format (*ID-q.tsv*) containing at least the quantified features together with their quantitative values. Any tsv table may be used as an *ID-q* file, provided that quantitative values are arranged in a table with column headers, so that features (e.g. PSMs or peptides) are arranged in rows and their quantitative values in columns, where every column pertains to a different sample. The column headers of the *ID-q* are used by iSanXoT to extract the necessary information.

In addition, when the *ID-q* file contains features quantitated in more than one experiment (e.g. different samples labelled with the same TMT-18plex tags), an additional column with the header *Experiment* must be included indicating the experiment ascription of the features.

Finally, iSanXoT also needs information to create the relation files required to integrate the quantified features into higher levels. This information is usually also present in the *ID-q* file. For instance, iSanXoT can use the columns containing the scan and the peptide identifiers to construct the *scan2peptide* relation table.

The majority of proteomics software tools generate tables that can be easily used for this purpose. In this Section we shall describe how to prepare the *ID-q* file based on the output from the three most popular proteomics pipelines (*Table S1*).

*Table S1. Output data from proteomics pipelines to be included in the* ID-q.tsv *file.*

| Proteomics pipeline | Experiment type | Output file name / suffix | Level name in the output file[1] | | | Quantitative data |
|---|---|---|---|---|---|---|
| | | | Scan | Peptide | Protein | |
| Proteome Discoverer (version 2.5) | Label-free | *_PeptideGroups.txt* | | *Sequence + Modifications* | *Master Protein Accessions* | *Abundance: FX: Sample Type* |
| | Isotopically labelled | *_PSMs.txt* | *Spectrum File + First Scan*[2] | *Sequence + Modifications* | *Master Protein Accessions* | *Abundance: Quan Channel* |
| MaxQuant (version 1.6.5.0) | Label-free | *modificationSpecificPeptides.txt* | | *Sequence + Modifications* | *Proteins* | *Intensity Experiment* |
| | Isotopically labelled | *msmsScans.txt* | *Raw file + Scan number* | *Modified Sequence* | *Proteins* | *Reporter intensity n* |
| Fragpipe (version 1.8.1) | Label-free | *combined_modified_peptide.tsv* | | *Modified Sequence* | *Protein ID* | *Experiment Intensity* |
| | Isotopically labelled | *psm.tsv* | *Spectrum + Spectrum File*[3] | *Modified Peptide* | *Protein ID* | *Channel* |

[1]If features quantitated in multiple experiments (e.g. different samples labelled with the same TMT-18plex tags) are to be considered, an additional column with the header *Experiment* must be included indicating the experiment ascription of the features.

[2]Make sure *Max. Number of Peptides Reported* = 1 was selected in the *Input Data* section of the Proteome Discoverer *Processing* node used.

[3]Make sure *Report top N* = 1 was selected in the *Advanced Output Options* of the FragPipe *MSFragger* module.

### Preparing the *ID-q* file from Proteome Discoverer output

In the case of Proteome Discoverer version 2.5 [8], the way that quantitative data are adapted for use with iSanXoT depends on whether they originate from label-free or labelled experiments:

#### Label-free experiments

In this case, quantitative data at the peptide level can be adapted for use with iSanXot from the *_PeptideGroups.txt* files obtained when the *Processing* workflow node *Minora Feature Detector* of Proteome Discoverer is used. The following column headers of the *_PeptideGroups.txt* files must be considered for preparing the *ID-q* file:

- *Sequence*: Amino acid sequence of the identified peptide;
- *Modifications*: Chemical or posttranslational modifications to the *Sequence* above;
- *Master Protein Accessions*: Accession code(s) for the protein(s) to which the peptide *Sequence* is ascribed;
- *Abundance: FX: Sample Type:* Peptide intensity in the RAW file identified with *FX* and tagged as *Sample Type* in the Proteome Discoverer *Input Files* tab.

The *peptide* level required for the *peptide to protein* integration with iSanXoT can be obtained by merging the *Sequence* and *Modifications fields* (see Section *Adapting the results from proteomics pipelines for iSanXoT* below).

#### Labelled experiments

For labelled experiments (e.g. TMT- or iTRAQ-based), quantitative data at the scan level can be adapted for use with iSanXot from the *_PSMs.txt* files generated when the *Processing* workflow node *Reporter Ions Quantifier* of Proteome Discoverer is used. The following column headers of the *_PSMs.txt* files must be considered for preparing the *ID-q* file:

- *Spectrum File*: Name of the RAW file where the PSM was identified;
- *First Scan*: Spectrum (scan) number of the PSM in the RAW file;
- *Sequence*: Amino acid sequence of the identified peptide;
- *Modifications*: Chemical or posttranslational modifications to the *Sequence* above;
- *Master Protein Accessions*: Accession code(s) for the protein(s) to which the peptide *Sequence* is ascribed;
- *Abundance: Quan Channel*: Intensity of the reporter ion tagged as *Quan Channel* in the Proteome Discoverer *Samples* tab.

For the *scan to peptide* integration with iSanXoT, the *scan* level can be obtained by merging the *Spectrum File* and *First Scan* fields, and the *peptide* level by merging the *Sequence* and *Modifications* fields (see Section *Adapting the results from proteomics pipelines for iSanXoT* below; make sure *Max. Number of Peptides Reported* = 1 was selected in the *Input Data* section of the Proteome Discoverer *Processing* node used).

### Preparing the *ID-q* file from MaxQuant output

The way MaxQuant version 1.6.5.0 [7] data are adapted for use with iSanXoT depends on whether they originate from label-free or labelled proteomics experiments:

#### Label-free experiments

In this case, the quantifications at the peptide level required to prepare the *ID-q* file can be found in the *modificationSpecificPeptides.txt* file, which is stored in the *"…combined/txt"* folder. The following column headers of the *modificationSpecificPeptides.txt* file must be considered for preparing the *ID-q* file:

- *Sequence*: Amino acid sequence of the identified peptide;
- *Modifications*: Chemical or posttranslational modifications to the *Sequence* above;
- *Proteins*: Identifier(s) of the protein(s) to which the peptide *Sequence* is ascribed;
- *Intensity Experiment*: Summed up extracted ion current of all isotopic clusters associated with the peptide Sequence identified across the raw files included in the *Experiment* as specified by the user in the MaxQuant *Raw data* tab.

The *peptide* level required for the *peptide* to protein integration with iSanXoT can be obtained by merging *Sequence* and *Modifications* fields (see Section *Adapting the results from proteomics pipelines for iSanXoT* below).

#### Labelled experiments

When dealing with labelled experiments (e.g. iTRAQ- or TMT-based), the necessary quantitative data at the scan level can be found in the *msmsScans.txt* file, which is stored in the *"…combined/txt"* folder. The following column headers of the *modificationSpecificPeptides.txt* file must be considered for preparing the *ID-q* file:

- *Raw file*: Name of the RAW file where the PSM was identified;
- *Scan number*: Spectrum (scan) number of the PSM in the RAW file;
- *Modified Sequence*: Amino acid sequence of the identified peptide including chemical or posttranslational modifications. This parameter is nonblank only when identification was successful.
- *Proteins*: Identifier(s) of the protein(s) to which the peptide *Sequence* is ascribed;
- *Reporter intensity n*: Intensity of the reporter ion *n* as specified by the user in the MaxQuant *Group-specific parameters* tab.

For the *scan to peptide* integration with iSanXoT, the *scan* level can be obtained by merging the *Raw File* and *Scan number* fields (see Section *Adapting the results from proteomics pipelines for iSanXoT* below).

### Preparing the *ID-q* file from FragPipe output

The way that quantitative data from Fragpipe version 1.8.1 [9] are adapted for use with iSanXoT depends on whether they originate from label-free or labelled experiments:

#### Label-free experiments

FragPipe *Quant (MS1)* module stores the quantifications at the peptide level necessary to prepare the *ID-q* file in a *combined_modified_peptide.tsv* file. The following column headers of the *modificationSpecificPeptides.txt* file must be considered for preparing the *ID-q* file:

- *Modified Sequence*: Amino acid sequence of the identified peptide;
- *Protein ID*: Identifier of the protein to which the *Modified Sequence* peptide is ascribed;
- *Experiment Intensity*: Summed up intensity of the *Modified Sequence* peptide in the RAW files included in the *Experiment* as specified by the user in the FragPipe *Workflow* tab.

#### Labelled experiments

Fragpipe *Quant (Isobaric)* module generates a *psm.tsv* output file that contains the quantitative data at the scan level obtained from labelled experiments. The following column headers of the *psm.txt* file must be considered for preparing the *ID-q* file:

- *Spectrum*: Spectrum (scan) identifier of the PSM in the XML file;
- *Spectrum File*: Name of the XML file where the PSM was identified;
- *Modified Peptide*: Amino acid sequence of the identified peptide including chemical or posttranslational modifications;
- *Protein ID:* Identifier of the protein to which the *Modified Peptide* is ascribed;
- *Channel*: Intensity of the reporter ion *Channel* as specified by the user in the FragPipe TMT-Integrator table of the *Quant (Isobaric)* module.

The *scan* level required for the later *scan to peptide* integration with iSanXoT can be obtained by merging the *Spectrum* and *Spectrum File* fields (see Section *Adapting the results from proteomics pipelines for iSanXoT* below; make sure *Report top N = 1* was selected in the *Advanced Output Options* of the FragPipe *MSFragger* module).

### Adapting the results from proteomics pipelines for iSanXoT

iSanXoT requires an identification/quantification tab-separated values file (*ID-q.tsv*) containing at least the identified features together with their quantitative values (an experiment identifier is required if two or more experiments are included). Users can either compose this *ID-q* file manually (see the previous Section to learn how to do that with data from the four most popular proteomics pipelines) or have it prepared by the iSanXoT Input Adaptor. The latter option is described in this Section.

- Run the iSanXoT application and create a new project or open an existing project. A new window will appear asking for the ID-q file (*Figure S43*).

- If you already have a suitable *ID-q* file, click the *Select User-Provided* option and then *Choose identification file* to select the file (*Figure S43*).

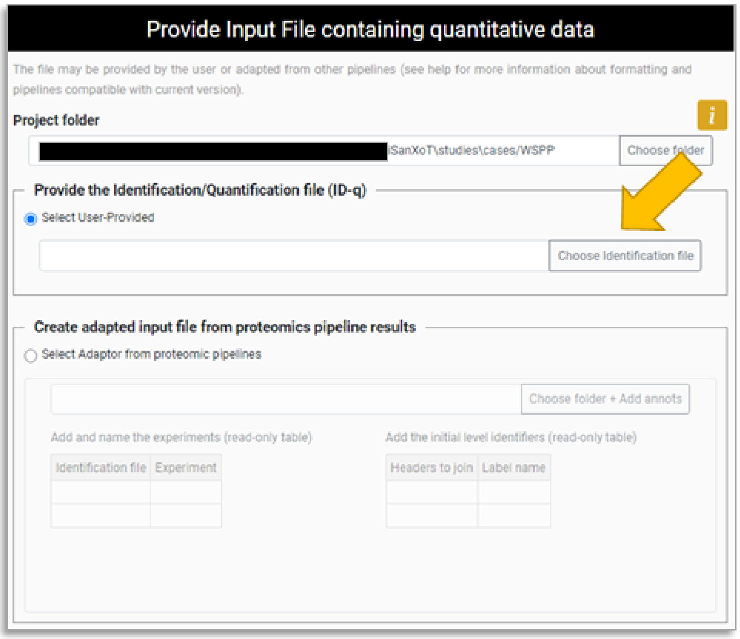

*Figure S43. Selecting an ID-q file in the Input Adaptor main window.*

- If you do not have an ID-q file, click *Select Adaptor from proteomics pipelines*. This option will launch the iSanXoT adapter to import your quantitative data. The adapter has been tested with recent versions of MaxQuant, Trans-Proteomic Pipeline, FragPipe, and Proteome Discoverer. Click on *Choose folder + Add anots* to select the folder containing your quantitative data (*Figure S44*). A three-panel window will pop-up (*Figure S45).*

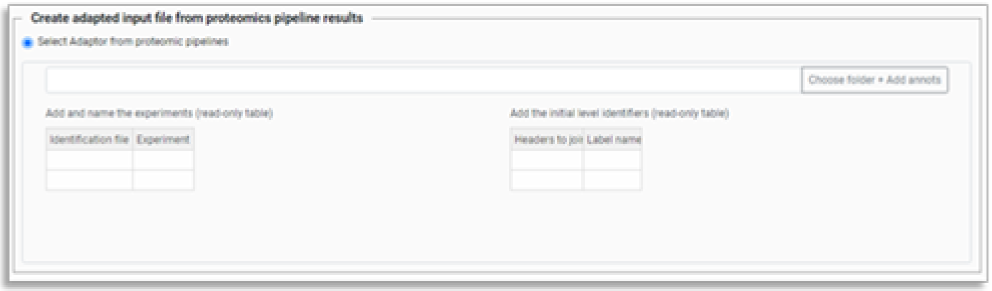

*Figure S44. Having the iSanXoT Input Adaptor prepare the ID-q file.*

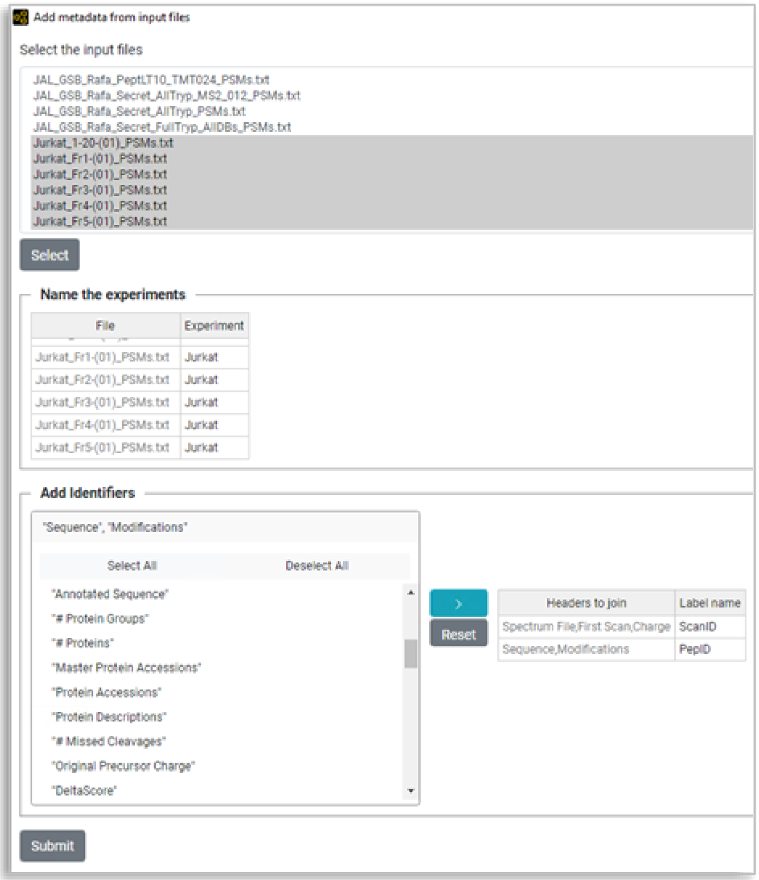

*Figure S45. Adapting results from a proteomics pipeline. In the top panel several output files from Proteome Discoverer have been selected. These PSMs.txt files, which contain identification/quantification data, have been assigned an experiment name (Jurkat) in the middle panel. The bottom panel has been used to create identifiers by concatenating result file headers: ScanID (by concatenating Spectrum File, First Scan and Charge) and pepID (by concatenating Sequence and Modifications).*

- The top panel displays the files included in the folder so that you can select one or more result files to be considered by the adapter. Please bear in mind that if several result files are selected, these must necessarily have the same column headers.
- The middle panel is used to set the distribution of data items across experiments according to result filenames.
- The bottom panel allows to create identifiers by concatenating result file headers. It is composed of two interfaces:
    - The headers found on the result files are listed on the left. Header names will be added to the interface on the right as they are selected;
    - The interface on the right displays the header names selected to generate the identifier as well as the identifier name provided by the user.

- Please note that the alphanumeric text that unambiguously identifies the items to be integrated is the only identifier that must be necessarily included in the *ID-q* file.
- Click the *Submit* button and the Input Adaptor will start generating the *ID-q.tsv* file.
